# Implementation of an antibody validation procedure: Application to the major ALS/FTD disease gene C9ORF72

**DOI:** 10.1101/499350

**Authors:** Carl Laflamme, Paul McKeever, Rahul Kumar, Julie Schwartz, Mahshad Kolahdouzan, Carol X.-Q. Chen, Zhipeng You, Faiza Benaliouad, Opher Gileadi, Heidi M. McBride, Thomas M. Durcan, Aled M. Edwards, Luke M. Healy, Janice Robertson, Peter S. McPherson

## Abstract

Antibodies are a key resource in biomedical research yet there are no community-accepted standards to rigorously characterize their quality. Here we develop a procedure to validate pre-existing antibodies. Cell lines with high expression of a target, determined through a proteomics database, are modified with CRISPR/Cas9 to knockout (KO) the corresponding gene. All commercial antibodies against the target are purchased and tested by immunoblot comparing parental and KO. Validated antibodies are used to definitively identify the most highly expressing cell lines, new KOs are generated if needed, and the lines are screened by immunoprecipitation and immunofluorescence. Selected antibodies are used for more intensive procedures such as immunohistochemistry. The pipeline is easy to implement and scalable. Application to the major ALS disease gene C9ORF72 identified high-quality antibodies revealing C9ORF72 localization to phagosomes/lysosomes. Antibodies that do not recognize C9ORF72 have been used in highly cited papers, raising concern over previously reported C9ORF72 properties.

## Introduction

Antibodies are vital tools in the biomedical research arsenal and in keeping with their importance there are over a million unique antibodies that are commercially available (Dr. Matt Baker, ThermoFisher, personal communication). In addition are the outputs of numerous, publicly-funded projects, which have generated recombinant or monoclonal antibodies, thus creating renewable antibodies with the potential for high reproducibility (Hornsby et al., 2015; Marcon et al., 2015; Na et al., 2016; Venkataraman et al., 2018; Andrews et al., 2019). The largest public antibody initiative has generated greater than 24,000 antibodies corresponding to almost 17,000 human genes, with a focus on polyclonal antibodies (Uhlen et al., 2015). Thus, there are pre-existing antibodies for a large percentage of the human genome.

The pace and attention given to antibody generation has not been matched with an equal effort on antibody characterization or in implementing standardized antibody characterization procedures. As a result, most products are generated and then sold/distributed with only rudimentary and non-quantitative supporting data, and as a result there are serious flaws in the reliability of many of the available reagents. Ill-defined antibodies contribute significantly to a lack of reproducibility in important research efforts including preclinical studies, with estimates that up to 90% of a select group of 53 landmark preclinical studies suffered from such flaws (Bradbury and Plückthun, 2015).

Several *ad hoc* international working groups have met to help define best practices for antibody validation (Taussig et al., 2018). One of the groups (Uhlén et al., 2016) proposed 5 separate validation criteria: 1) genetic strategies in which the specificity of the antibody toward the endogenous protein is confirmed by the loss of signal in cells or tissues comparing parental to knockout (KO) or knockdown controls; 2) orthogonal strategies in which correlations are made between the antibody signal and known information regarding protein abundance or localization; 3) the ability of two independent uncharacterized antibodies recognizing different epitopes in the same target protein to recognize the same protein; 4) using overexpressed epitope-tagged proteins comparing antibodies against the tag to the uncharacterized antibody; 5) immunoprecipitation followed by mass spectrometry to determine if the protein of interest is a major signal in the sample. These criteria are arguably not of equal scientific value and there is no consensus as to which should be used. The first and fifth methods are the most unbiased and useful, whereas the remaining are less informative and perhaps flawed. The genetic approaches presented in criterion 1 are suitable for antibody validation in all applications, yet there is no template for their application.

Historically, the lack of a suitable control – an isogenic source of proteins lacking the target antigen, has hampered the implementation of criterion 1, but this has changed: it is now routine to make KO cell lines in an array of cell types which, for non-essential proteins, provides the ideal control for testing antibody specificity for the endogenous protein in multiple applications. This capability then opens up the possibility of creating a standardized characterization process that can be applied systematically to characterize not only new antibodies but also the ~1 million antibodies that are already commercially available. With such a process in hand it should be possible to identify high quality antibodies for different applications from the existing set of commercially-available antibodies, seemingly for a large percentage of all human gene products.

To pilot the concept that excellent antibodies can be found among those that are commercially available if one carries out a systematic analysis, we studied the major amyotrophic lateral sclerosis (ALS, OMIM #105400) disease gene product, C9ORF72. ALS is a fatal neurodegenerative disease characterized by progressive paralysis leading to respiratory failure (Kiernan et al., 2011) and is on a genetic and pathophysiological continuum with frontotemporal dementia (FTD, OMIM #600274) (Ng et al., 2015). A search for genes involved in ALS/FTD led to the discovery of a hexanucleotide-repeat expansion mutation in the first intron of *C9ORF72*. In a North America cohort, this mutation underlies 11.7% and 23.5% of familial FTD and ALS cases, respectively, and in a large Finnish population, the *C9ORF72* mutation underlies 46.0% of familial ALS and 21.1% of sporadic ALS (DeJesus-Hernandez et al., 2011; Renton et al., 2011). Thus, the *C9ORF72* mutation is the most common genetic abnormality in both FTD and ALS. It is vital to understand the cell biological role of C9ORF72, but the literature regarding C9ORF72 subcellular and tissue-distribution is conflicting (Burk et al., 2019). We believe the lack of consensus on C9ORF72 expression, function and subcellular localization stems in part from the use of non-specific antibodies.

C9ORF72 provided an excellent protein on which to develop an antibody characterization process because although the protein is of relatively low abundance, there are many commercially-available antibodies. The process we outline can be applied to any protein target and here it led us to the recognition of problems with the C9ORF72 literature and to the discovery of unexpected properties of the protein.

## Results

### Development of an antibody validation strategy

The antibody validation strategy developed in this manuscript is presented in overview in Figure 1. The steps were built empirically as we proceeded with our analysis of antibodies for C9ORF72. The workflow is as follows: 1) use PaxDB (https://pax-db.org/) to identify a cell line that expresses the protein of interest at relatively high levels, is readily modifiable by CRISPR/Cas9, and is easy to grow and manipulate; 2) use CRISPR/Cas9 to generate a KO in this cell line; 3) identify all commercial antibodies against the protein of interest and screen them by immunoblot using parental and KO controls; 4) use an antibody validated in step 3 for quantitative immunoblots on a panel of cell lines to identify a line that has the highest levels of expression of the target protein and has appropriate characteristics for the target of interest. This step is recommended as it provides more reliable information than could be provided by the proteomics database. However, the selected line may well be the original line that was chosen; 5) use the selected edited line to screen antibodies for specificity by immunoprecipitation and immunofluorescence.

**Figure 1:**
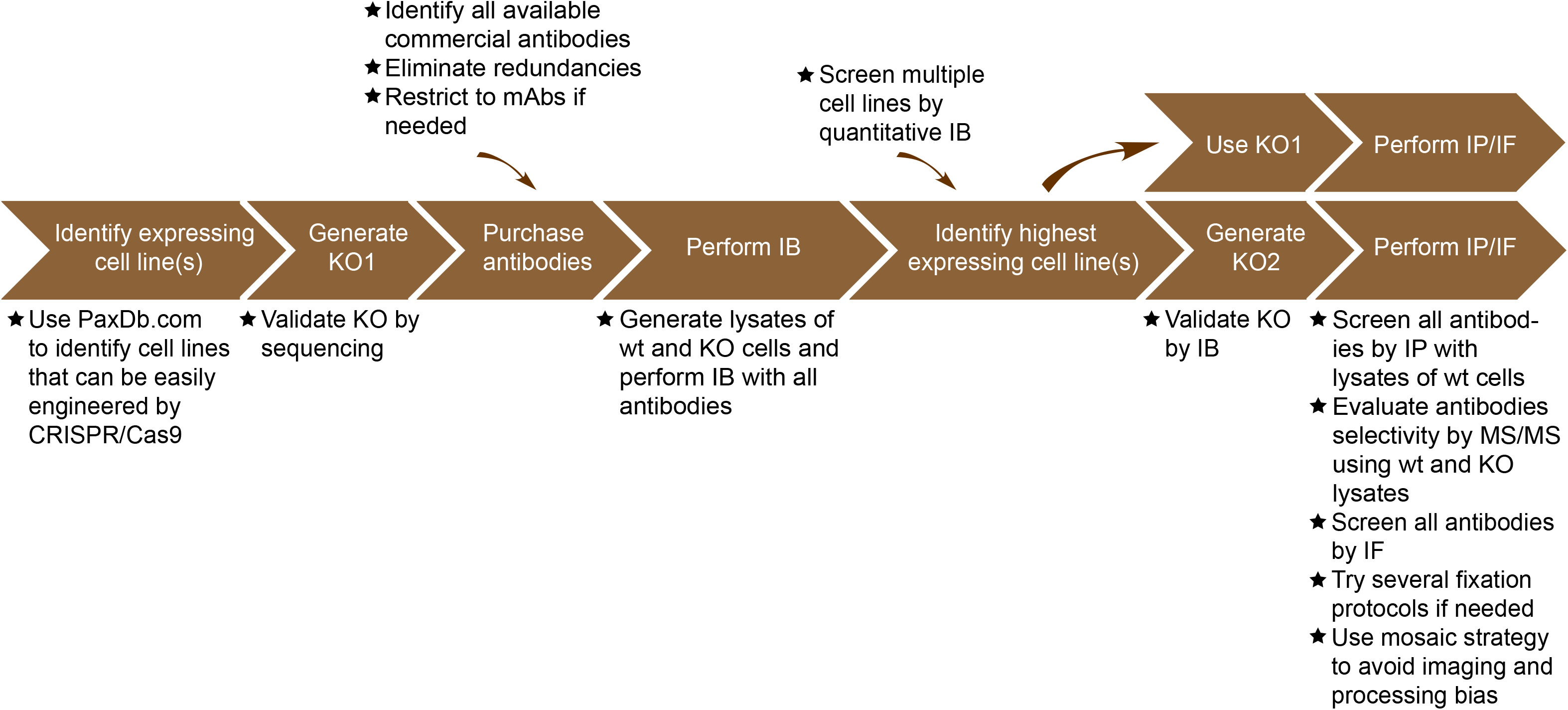
Summary of the antibody validation pipeline. IB=immunoblot; IP=immunoprecipitation; IF=immunofluorescence

### Selection and editing of a cell line and selection of antibodies

The single most common application of commercial antibodies for the majority of laboratories is immunoblot. We initially sought to screen all commercially available C9ORF72 antibodies by immunoblot using parental and KO cell lines. We first referenced PaxDb, a comprehensive proteomic database that provides protein abundance across multiple organisms and tissues/cell lines (Wang et al., 2015) (https://pax-db.org/). C9ORF72 is expressed at relatively low levels in most tissues and cell lines ranging from the 77^th^ percentile for HeLa cells to the 35^th^ percentile in RKO cells. We eventually chose to start with HEK-293 cells (expression levels at the 65^th^ percentile) as they are the 5^th^ most abundant C9ORF72 expressing cell line, and are easy to grow and to edit with CRISPR/Cas9. We then used CRISPR/Cas9 to generate C9ORF72 heterozygous and KO HEK-293 cells. Briefly, we introduced the sequence of our KO sgRNA into an expression plasmid bearing both the sgRNA scaffold backbone and Cas9. We then transfected the plasmid into HEK-293 cells and selected transfected cells with puromycin for 3 days. Cells were then grown following clonal dilution and sequencing of the genomic DNA of multiple clones revealed double-strand breaks at the expected localization leading to one heterozygous and two complete KO lines. The complete process took ~5 weeks. We then purchased all known commercial antibodies for C9ORF72. For this we searched the literature by PubMed and examined the websites of companies with C9ORF72 antibodies identified using Google searches. We initially identified more than 100 antibodies for C9ORF72. However, we avoided purchasing the same antibody available from different companies. For example, the C9ORF72 antibody *Prestige Antibodies® Powered by Atlas Antibodies* was purchased at Sigma (HPA023873), but is also sold by other companies including Novus Biological (NBP1-93504). This led to 12 distinct commercial C9ORF72 antibodies, 9 rabbit polyclonal antibodies and 3 mouse monoclonal antibodies (Table 1). The antibodies are available from Abcam (4), Proteintech (3), GeneTex (3), Sigma (1), and Santa Cruz (1). In addition we identified 2 sheep polyclonal antibodies from the Protein Phosphorylation and Ubiquitylation Unit of the Medical Research Council (MRC) in Dundee Scotland (Table 1). Interestingly, we eventually received refunds for many of the antibodies that did not perform as advertised, lowering overall costs.

**Table 1.** Summary of C9ORF72 antibodies and their properties

| Companies | Catalog number | Used in # papers | Citations to papers | Vendors proposed application(s) | Epitope | Host/clonality | Wb | IF | IP |
| --- | --- | --- | --- | --- | --- | --- | --- | --- | --- |
| Abcam | ab227555 | 0 | 0 | Wb/IF-ICC/IHC | a.a.395-424 | Rabbit polyclonal | - | - | + |
|  | ab171428 | 0 | 0 | Wb | Sequence is proprietary | Rabbit polyclonal | - | - | - |
|  | ab121779 | 1 | 63 | IF-ICC/IHC | a.a.110-199 | Rabbit polyclonal | + | - | + |
|  | ab203627 | 0 | 0 | IHC | a.a.390-440 | Rabbit polyclonal | - | - | - |
| Proteintech | PT25757 | 1 | 16 | Wb/IHC/IF | a.a. 1-221 | Rabbit polyclonal | + | - | + |
|  | PT22637 | 7 | 164 | Wb/IHC/IF | a.a. 1-169 | Rabbit polyclonal | + | - | + |
|  | PT66140 | 0 | 0 | Wb/IHC/IF | a.a. 1-222 | mouse monoclonal | - | - | - |
| GeneTex | GTX632041 | 0 | 0 | Wb/IHC/IP | Sequence is proprietary | mouse monoclonal | ++ | +++ | +++ |
|  | GTX634482 | 0 | 0 | Wb | Sequence is proprietary | mouse monoclonal | +++ | - | - |
|  | GTX119776 | 5 | 757 | Wb/IHC/IP/IF | Sequence is proprietary | Rabbit polyclonal | - | - | + |
| Sigma | HPA023873 | 10 | 1091 | IHC | a.a.110-199 | Rabbit polyclonal | + | - | + |
| Santa Cruz | sc-138763 | 15 | 3164 | Wb/IHC/IP | a.a.165-215 | Rabbit polyclonal | - | - | - |
| MRC | MRC-S478D | 0 | 0 | Wb/IP | full length | Sheep polyclonal | ++ | - | + |
|  | MRC-S479D | 0 | 0 | Wb/IP | full length | Sheep polyclonal | + | - | + |

### Analysis by immunoblot

We screened the 14 C9ORF72 antibodies by immunoblot using lysates of HEK-293 cells including parental cells, the isogenic heterozygous line and the two isogenic KO lines (Fig. 2). For each series of blots we resolved 50 μg of protein (using 5-16% gradients gels) from lysates prepared in buffer containing 1% Triton X-100 in order to extract both cytosolic and membrane-associated proteins. The proteins, transferred to nitrocellulose membranes, were stained with Ponceau to ensure even loading of lysates. After screening all 14 antibodies we found one mouse monoclonal antibody from GeneTex (GTX634482) that has optimal features. It shows a strong signal in parental HEK-293 cell lysates, the signal is reduced by ~50% in the heterozygous line, and no signal for the antibody is detected in the two KO lines. Moreover, there are no other bands visible (Fig. 2), even upon long exposure (Fig. S1). Thus, the signal to noise ratio in HEK-293 cells is sufficient to justify their use in immunoblot screens. GeneTex monoclonal GTX632041 also recognizes C9ORF72 seemingly selectively in the short exposure (Fig. 2), but other bands are detected following longer exposure (Fig. S1). Other antibodies that recognize C9ORF72 include ab12179 from Abcam, PT25757 and PT22637 from Proteintech, HPA023873 from Sigma and MRC-S478D and MRC-S479D from the MRC. However, all of these antibodies detect additional bands, and in several cases they have multiple cross-reactivity. Six of the antibodies do not detect endogenous C9ORF72. Thus, GTX634482 from GeneTex is a robust antibody for immunoblot.

**Figure 2:**
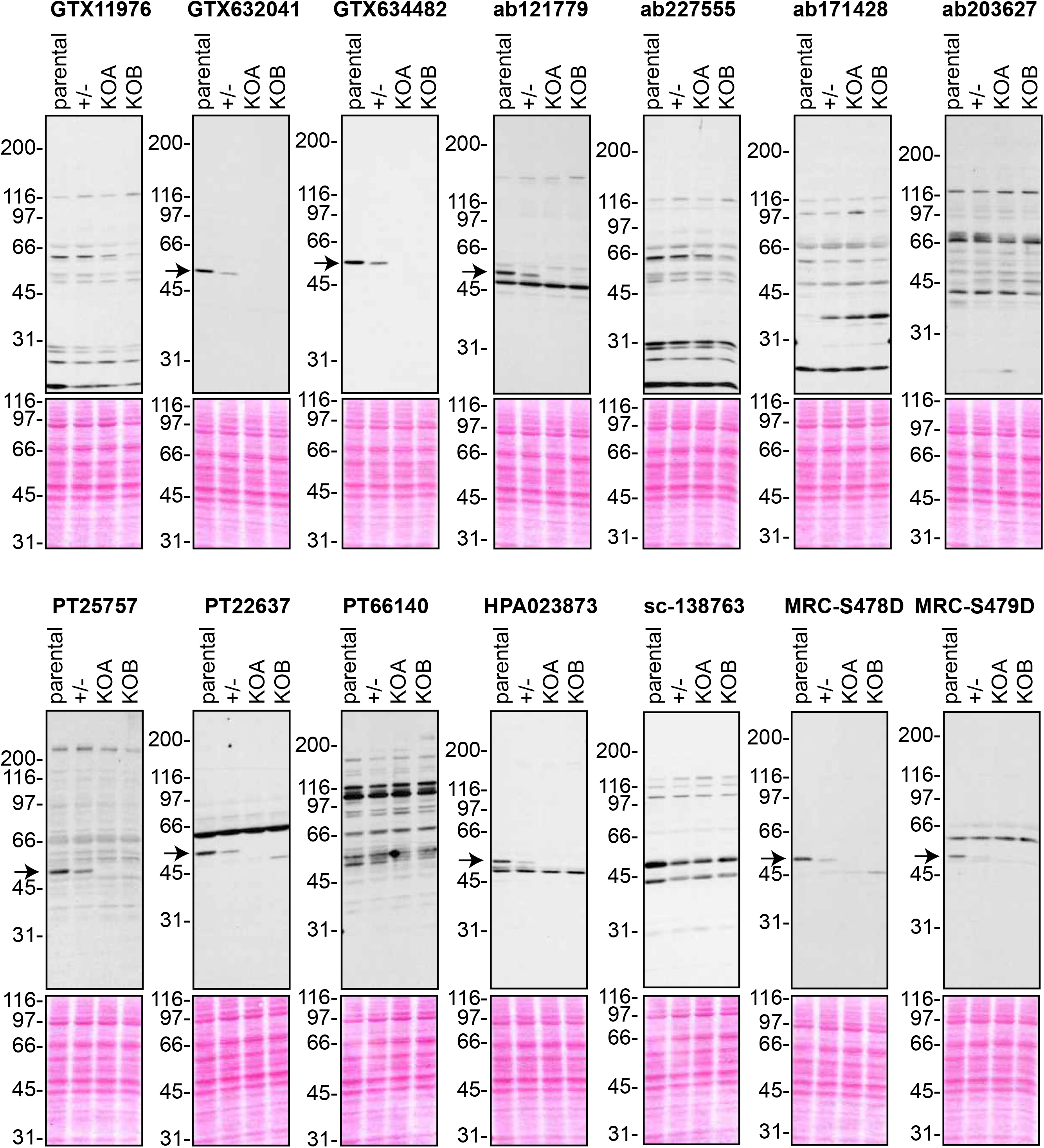
Analysis of C9ORF72 antibodies by immunoblot. Cell lysates from HEK-293 parental, heterozygous (+/-) or two individual C9ORF72 KO clones (KOA, KOB) were prepared and processed for immunoblot with the indicated C9ORF72 antibodies. The arrows point to positive C9ORF72 signals. The Ponceau stained transfers associated with each blot are shown as a protein loading control.

### Analysis by immunoprecipitation

We next sought to test the functionality of the antibodies in immunoprecipitation applications. All 14 antibodies were pre-coupled to protein A- or protein G-Sepharose as appropriate and a detergent solubilized HEK-293 cell lysate was prepared and incubated with the antibody/bead mixtures. For controls lysates were incubated with beads alone or the bead/antibody conjugates were incubated with buffer alone. After washing the beads, the presence of C9ORF72 in the immunoprecipitates was detected by immunoblot using rabbit antibody PT22637 for the immunoprecipitates with mouse antibodies and the mouse antibody GTX634482 for immunoprecipitates with rabbit or sheep antibodies (Fig. 3). Of the 14 antibodies, 9 immunoprecipitate endogenous C9ORF72, with GeneTex monoclonal antibody GTX632041 demonstrating the most robust enrichment of the protein compared to starting material (Fig. 3A, 3B). Five antibodies showed no appreciable C9ORF72 immunoprecipitation including GTX634482 (Fig. 3A, 3B), which was the most effective antibody in immunoblot. The effectiveness of GTX634482 on immunoblot coupled to its inability to immunoprecipitate C9ORF72 highlights the need to screen all antibodies for all applications.

**Figure 3:**
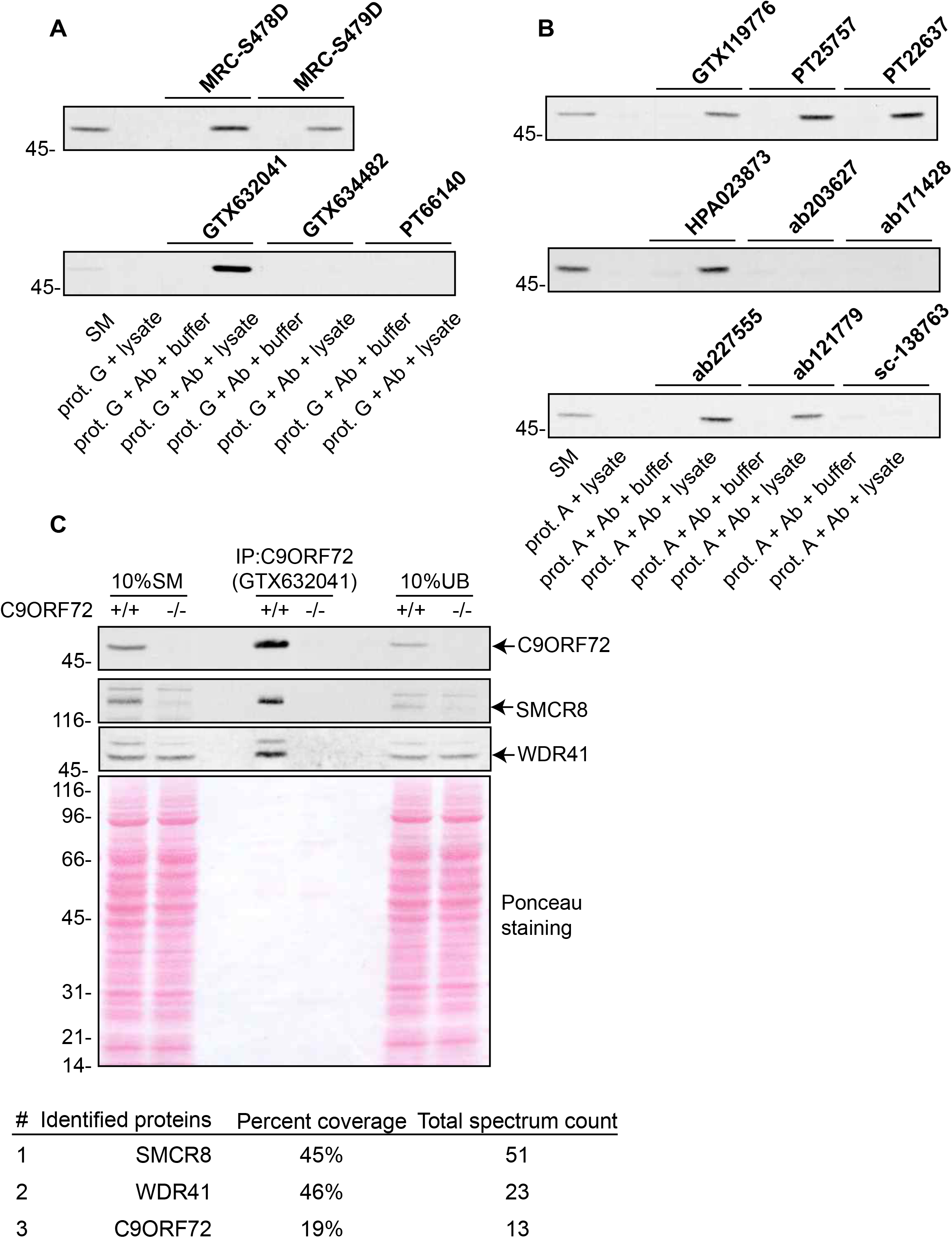
Analysis of C9ORF72 antibody by immunoprecipitation. **(A)** HEK-293 cell lysates were prepared and immunoprecipitation was performed using the indicated C9ORF72 monoclonal antibodies pre-coupled to protein G-Sepharose (prot. G). Controls included the Protein G alone incubated with cell lysate or protein G beads pre-coupled with the antibodies but incubated with lysis buffer. Samples were washed and processed for immunoblot with C9ORF72 antibody PT22637. **(B)** HEK-293 cell lysates were prepared and immunoprecipitation was performed using the indicated C9ORF72 polyclonal antibodies pre-coupled to protein A-Sepharose (prot. A). Controls were as in **A**. Samples were washed and processed for immunoblot with C9ORF72 antibody GTX634482. **(C)** Lysates were prepared from HEK-293 cells, parental (+/+) and KO (-/-) and immunoprecipitation was performed using C9ORF72 antibody GTX632041 pre-coupled to protein G-Sepharose. Samples were washed and processed for immunoblot with C9ORF72 antibody PT22637. Blots were also performed for SMCR8 and WDR41. The Ponceau stained transfer of the blot is shown as a protein loading control. Aliquots (10%) of the cell lysates before (starting material, SM) and after incubation with the antibody-coupled beads (unbound, UB) were processed in parallel. The table shows the 3 top hits (total spectrum counts) obtained by mass spectrometry analysis of the immunoprecipitated samples.

Using mass spectrometry we assessed the degree to which GTX632041 was selective for C9ORF72 and able to co-purify SMCR8, a known obligate interacting partner of the protein (Sellier et al., 2016; Amick et al., 2016; Zhang et al., 2018). We performed immunoprecipitation studies with GTX632041 using parental and KO HEK-293 cell lysates. When we examined the soluble cell lysate (starting material, SM) and the unbound proteins (UB) of immunoprecipitations from the parental cells we observed that a significant proportion of C9ORF72 and SMCR8 are removed from the lysate by immunoprecipitation with antibody GTX632041 and that both proteins are strongly enriched in the immunoprecipitated sample relative to SM (Fig. 3C). Mass spectrometry of the samples immunoprecipitated with GTX632041 revealed C9ORF72 (13 peptides), SMCR8 (51 peptides), and another known C9ORF72-binding partner, WDR41 (23 peptides) (Sellier et al., 2016) (Fig. 3C). None of the other 13 antibodies allow for an appreciable co-immunoprecipitation of SMCR8 (data not shown). The presence of C9ORF72 and its known-binding partners at near 1:1:1 level (when adjusting for the mass of the proteins) further supports the use of HEK-293 cells in our analysis. No peptides are detected for any of these proteins by mass spectrometry of immunoprecipitates performed under identical conditions from lysates of C9ORF72 KO cells, further demonstrating the high selectivity of the antibody for C9ORF72.

### Analysis by immunofluorescence

We next tested the effectiveness of the antibodies for immunofluorescence applications. HEK-293 cells, parental and KO were fixed in paraformaldehyde (PFA), permeabilized with Triton X-100, and stained with C9ORF72 antibodies with appropriate fluorescent secondary antibodies. The antibodies gave various staining patterns with no reduction in fluorescent staining in KO cells (Fig. S2). Similar results were seen when the cells were fixed in −20°C methanol (Fig. S3).

Achieving successful detection of endogenous proteins by immunofluorescence can be challenging, especially for lower abundance proteins. In an effort to obtain a better signal to noise ratio, we sought to identify a cell line with higher levels of endogenous C9ORF72 expression. We thus used GTX634482, validated as described (Fig. 2) to determine the relative levels of C9ORF72 in multiple cell lines employing the LI-COR Odyssey Imaging System (LI-COR Biosciences), which utilizes fluorescent secondary antibodies to allow for quantitative immunoblots. C9ORF72 is detected in a variety of human cell lines including U2OS (osteocarcoma), HeLa (cervical cancer), RKO (colon carcinoma), U87 and U251 (glioblastomas), motor neuron precursor cells (NPCs) and motor neurons (MNs) derived from human-induced pluripotent stem cells (Fig. 4A). Of the cell lines tested, U2OS has the highest levels of C9ORF72 whereas RKO cells have lower expression than HeLa (Fig. 4A). Interesting, the levels of C9ORF72 in motor neurons is similar to that in HEK-293 (Fig. 4A). U2OS are easy to genome edit and are large cells suitable for immunofluorescence analysis. We thus used CRISPR/Cas9 to generate U2OS cells lacking C9ORF72, which was confirmed by immunoblot with GTX634482 (Fig. 4B). To better distinguish between the parental and C9ORF72 KO lines, we transfected the lysosomal protein LAMP1 tagged to YFP in parental cells and LAMP1-RFP in C9ORF72 KO cells. We then re-plated the cells such that both parental and KO cells were found on the same coverslip as a mosaic (Fig. 4C). This strategy reduces microscopy imaging and figure processing biases. Using this strategy we found GTX632041 as the sole antibody to recognize endogenous C9ORF72 specifically. Indeed, GTX632041 immunofluorescence labeling reveals a cytosolic/punctate fluorescence signal that is dramatically reduced in KO cells to a level comparable to buffer control (Fig. 4D). The other 13 antibodies showed non-specific staining patterns that are similar between control and KO cells (Fig. 4D and Fig. S4).

**Figure 4:**
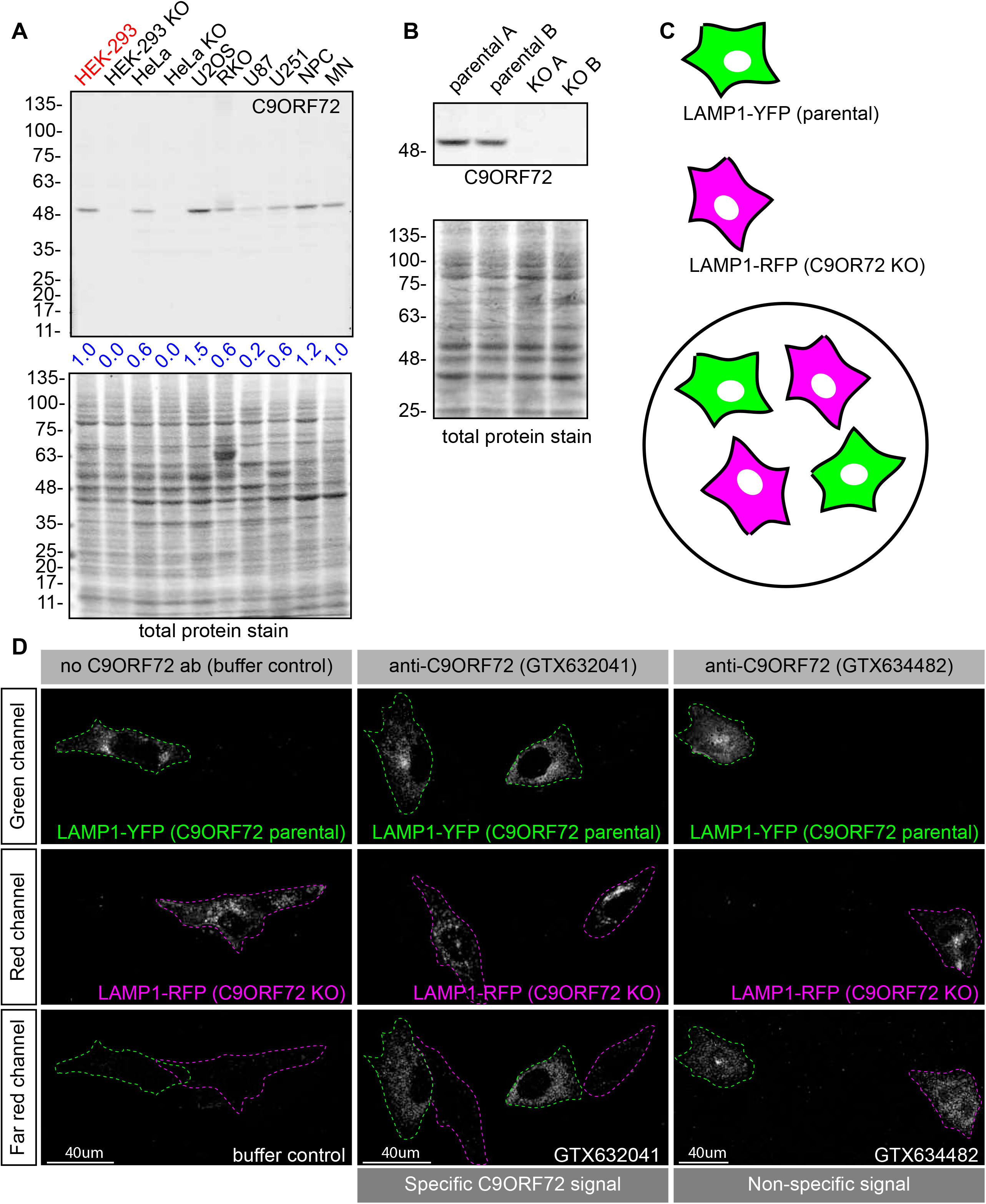
Analysis GeneTex C9ORF72 monoclonal antibodies by immunofluorescence. **(A)** Lysates were prepared from selective human cell lines and processed for immunoblot using antibody GTX634482 with the LI-COR Odyssey Imaging System that utilizes fluorescent secondary antibodies to allow for quantitative blots (HEK, human embryonic kidney; HeLa, human cervical cancer; U2OS, human osteocarcoma; RKO, human colon carcinoma; U87/U251, human glioblastoma; NPC, human neural precursor cells; MN, human motor neurons). The total protein stained transfers are shown as a loading control. The C9ORF72 protein signal as a ratio to total protein was determined, normalized to parental HEK-293 cells (red), and presented as fold change (blue numbers). **(B)** Cell lysates of parental and KO U2OS cells were analyzed by immunoblot with C9ORF72 antibody GTX634482. **(C)** Mosaic strategy used to investigate antibodies specificity by immunofluorescence. **(D)** Parental and KO cells were transfected with LAMP1-YFP or LAMP1-RFP, respectively. Parental and KO cells were combined and incubated with buffer only or stained with the indicated GeneTex C9ORF72 monoclonal antibodies. Greyscale images of the green, red and far-red channels are shown. Parental and KO cells are outlined with green and red dashed line, respectively. Representative images are shown. Bars = 40μm.

Based on this experience we recommend that once immunoblot screens identify a specific antibody for a target of interest, the next step should involve screening panels of cell lines with that antibody to find those with the highest expression levels (Fig. 1). While PaxDb appears to be effective to identify appropriate lines to start the analysis it is not infallible. For example, while it indicated higher expression levels for C9ORF72 in U2OS versus HEK-293, it also indicated that RKO had higher levels than U2OS, which did not bear out in the immunoblot analysis (Fig. 4A). Thus, moving straight to KO in U2OS cells for the immunofluorescence analysis would have been a more effective strategy. Moreover, while the HEK-293 cells were effective for the immunoprecipitation studies here (Fig. 3), we recommend also performing the immunoprecipitation studies following the cell distribution blots (Fig. 1).

### Identification of C9ORF72 antibodies effective for immunohistochemistry

Given the importance of C9ORF72 in ALS and FTD it is vital to identify antibodies effective for staining of tissue sections. We thus used diaminobenzidine (DAB) labeling comparing brain sections from wild-type littermate mice to mice with the C9ORF72 ortholog (31100432021Rik) deleted (Jiang et al., 2016). We originally focused on GTX634482, which is the strongest and most specific antibody for immunoblot (Fig. 2), and also recognizes the protein in mouse brain (Fig. S5). We tested the antibody on sections that had been treated at 110°C, pH 9.0 for epitope unmasking. A punctate and/or neuritic-like signal is observed in the neuropil of the glomerular layer of the olfactory bulb, the ventral pallidum of the basal ganglia, the CA4 (hilus), CA3, and CA2 region of the hippocampus, the substantia nigra, the inferior olive, and granular layer of the cerebellar cortex of wild-type mice (Fig. 5A, arrows). The signal is nearly completely ablated in the KO mice although there is some non-specific signal seen in the dentate gyrus/CA4, inferior olive and cerebellar cortex sections (Fig. 5A, arrowheads). The staining pattern observed is similar to the basal ganglia, hippocampal formation, and cerebellum staining observed in a previous study (Frick et al., 2018). GTX632041, which is the most effective antibody for immunoprecipitation (Fig. 3) and immunofluorescence (Fig. 4D) analysis, also specifically recognizes C9ORF72 in mouse brain sections including those through the hippocampus (Fig. 5B) but staining is not as strong or consistently specific as seen with GTX634482.

**Figure 5:**
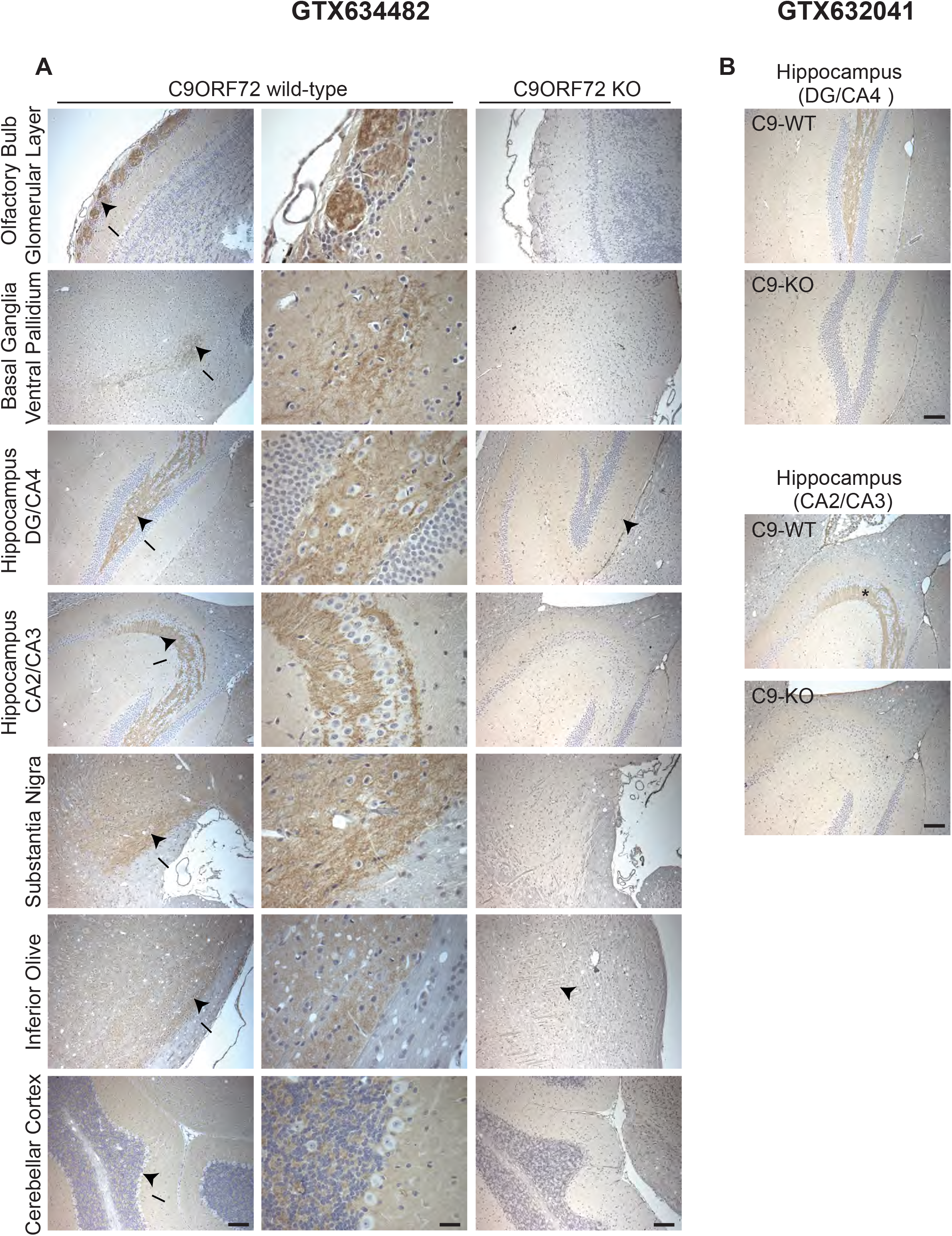
Analysis of C9ORF72 antibodies by immunohistochemistry. **(A)** DAB immunohistochemistry of C9ORF72 wild-type and C9ORF72 KO mouse brains in the sagittal plane using C9ORF72 antibody GTX634482. Micrographs are ordered from the rostral to caudal aspect of the mouse brain. For C9ORF72 WT arrows indicate areas of increased DAB labeling intensity and correspond to the inset regions on the right. Arrowheads indicate areas of non-specific labelling in the C9ORF72 KO mice. Scale bars = 100 μm or 25 μm for the inset images on the right for the C9ORF72 wild-type micrographs). **(B)** DAB immunohistochemistry of C9ORF72 wild-type and C9ORF72 KO mouse brains in the sagittal plane using C9ORF72 antibody GTX632041. Scale bars = 100 μm.

Thus, we have identified two mouse monoclonal antibodies that effectively and specifically recognize C9ORF72 in multiple applications. The antibody GTX634482 is recommended for immunoblot and immunohistochemical applications on antigen unmasked samples, and GTX632041 is recommended for immunoprecipitation and immunofluorescence.

### C9ORF72 protein levels peak in macrophages

The GeneTex antibodies GTX634482 and GTX632041 are valuable and powerful tools to assess C9ORF72 biology. We thus used these antibodies to resolve controversies regarding C9ORF72 tissue distribution and subcellular localization (Burk et al., 2019). We confirmed the utility of the anti-human GTX634482 in immunoblots of extracts from mouse tissues allowing us to use KO mice for control in tissue distribution blots. C9ORF72 is detected at the highest levels in mouse brain with lower levels in spinal cord, testis and the spleen (Fig. S5), as described previously (Frick et al., 2018). Lower levels of C9ORF72 protein are detected in a series of non-neuronal tissues and cell lines and no signal is observed in brains of C9ORF72 KO mice (Fig. S5).

C9ORF72 KO mice age without motor neuron disease (O’Rourke et al., 2016; Jiang et al., 2016; Burberry et al., 2016) but instead develop progressive splenomegaly and lymphadenopathy and ultimately die due to complications of autoimmunity (O’Rourke et al., 2016; Burberry et al., 2016; Atanasio et al., 2016). C9ORF72 is required for the normal function of murine myeloid cells, such as macrophages and microglia (resident macrophages in the central nervous system) (O’Rourke et al., 2016; Atanasio et al., 2016; Jiang et al., 2016) and C9ORF72 KO macrophages and microglia display lysosomal dysfunction (O’Rourke et al., 2016; Zhang et al., 2018). Interestingly C9ORF72 is detected in RAW264.7 cells, a mouse macrophage-like cell line (Fig. S5). We thus investigated C9ORF72 protein levels in primary myeloid cells compared to primary neurons (Fig. 6). We isolated murine bone marrow-derived macrophages (BMDM) and rat cortical and hippocampal neurons and compared C9ORF72 expression to whole mouse brain and to HEK-293 cells by quantitative immunoblot. Cortical and hippocampal neurons express C9ORF72 to an intermediate level between HEK-293 and murine whole brain (Fig. 6A). To our surprise, isolated BMDM have C9ORF72 protein levels that are 5.5-fold higher than HEK-293 cells, and 1.5-fold higher than whole brain (Fig. 5A).

**Figure 6:**
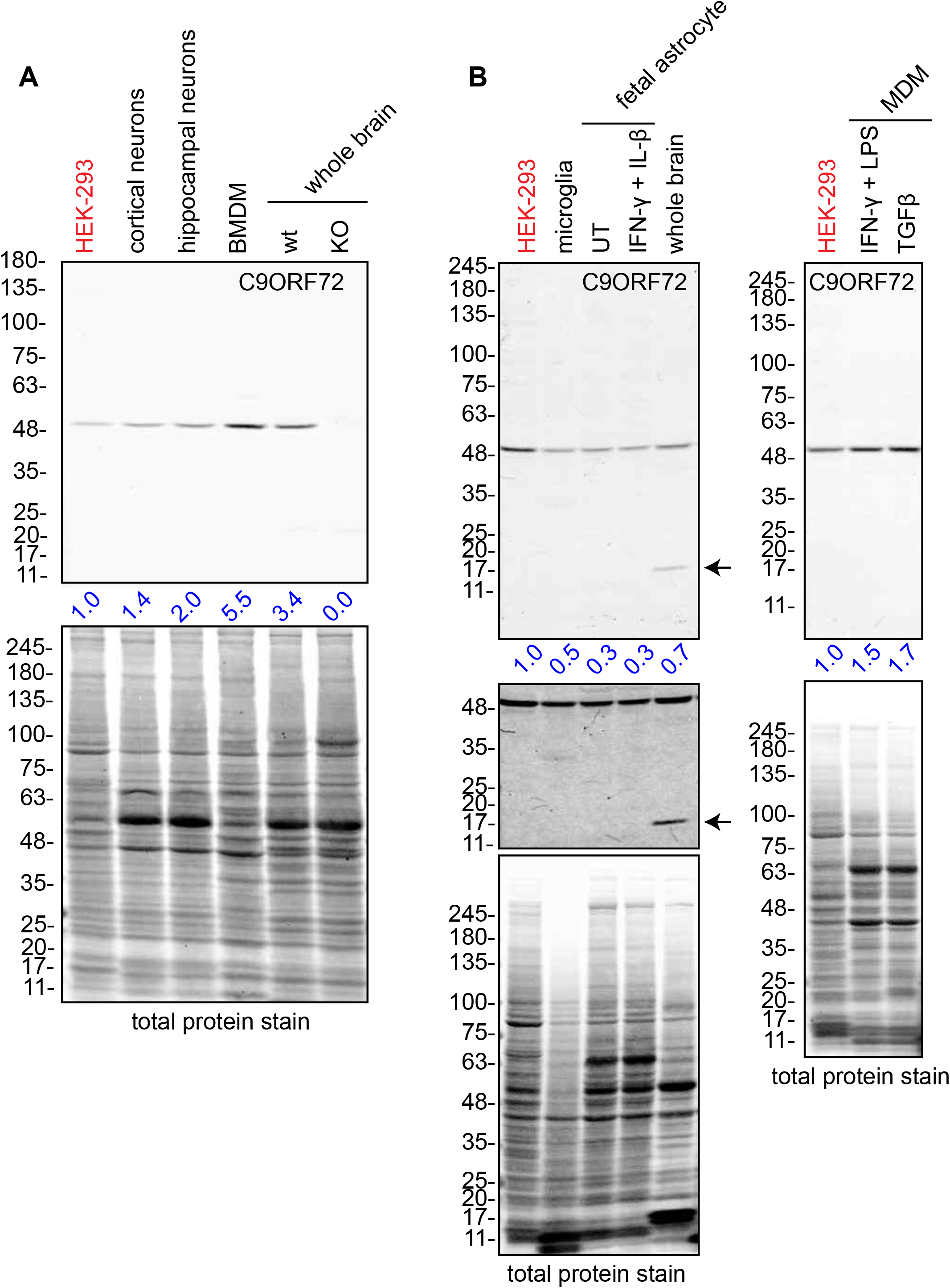
Murine and human macrophages express C9ORF72 at high levels. **(A)** Lysates were prepared from rat cortical neurons, rat hippocampal neurons, mouse bone marrow-derived macrophages (BMDMs) and mouse whole brain, and processed for quantitative immunoblot using antibody GTX634482. The total protein stained transfers are shown as a loading control. The C9ORF72 protein signal as a ratio to total protein was determined, normalized to parental HEK-293 cells (red), and presented as fold change (blue numbers). C9ORF72 KO mouse brain (KO) was included as a specificity control. **(B)** Lysates were prepared from human primary cells including microglia, fetal astrocytes either untreated (UT) or treated with IFN-γ combined with IL-β, human whole brain and monocyte-derived macrophages (MDMs) treated with either IFN-γ and LPS or with TGF-β. Immunoblot and C9ORF72 quantification were performed as in **A**.

We next investigated C9ORF72 protein levels in primary human cells. Isolated adult microglia, together with fetal astrocytes, activated or not with interleukin-β (IL-β) had lower C9ORF72 protein levels than HEK-293 cells (Fig. 6B). Human whole brain also had lower C9ORF72 protein expression than HEK-293 cells. Interestingly, we identified a 17 KDa protein that is reactive with GTX634482 and is expressed specifically in human samples (Fig. 6B). This protein is likely the predicted C9ORF72 short isoform, which has a theoretical molecular weight of ~25 kDa (Xiao et al, 2015). Next, we examined human monocyte-derived macrophages under traditional M1 polarization conditions (IFN-γ and LPS treated) or in the presence of TGF-β, which reproduces the homeostatic state of the cells *in vitro* (Healy et al., 2016) (Fig. 6B). We observed higher C9ORF72 expression in human macrophages under M1 or resting state than in HEK-293 cells (1.5 to 1.7 fold, respectively). Thus, C9ORF72 is expressed at the high levels in macrophages.

### C9ORF72 localizes to lysosomes and phagosomes

Endogenous C9ORF72 has been localized to the nucleus (Renton et al., 2011, Atkinson et al., 2015), early endosomes (Farg et al., 2014, Shi et al., 2018), recycling and late endosomes (Farg et al., 2014), the Golgi apparatus (Aoki et al., 2017), stress granules (Chitiprolu et al., 2018) the cytoplasm, neurites, growth cones and neuropils (Atkinson et al., 2015), and lysosomes (Shi et al., 2018). Each of these studies performed immunofluorescence using C9ORF72 commercial antibodies that failed our validation criteria.

In contrast, one study used CRISPR to edit endogenous C9ORF72 protein with an HA tag and used an antibody against the tag. This study revealed a diffuse cytoplasmic localization for tagged C9ORF72 under basal cell culture conditions, with the observation that the protein became highly concentrated on lysosomes following starvation (Amick et al., 2016).

To investigate the localization of endogenous C9ORF72 we used U2OS cells transfected with LAMP1-RFP and stained with C9ORF72 antibody GTX632041. Cells were examined using confocal microscopy followed by image deconvolution (HyVolution-Leica), either under basal conditions or following serum-starvation for 2 h. HyVolution increased image resolution (Fig. S6). Under basal conditions, C9ORF72 demonstrated a discrete punctate pattern with peripheral lysosomal localization (Fig. 7, top panel). In fact, 82% of LAMP1-positive lysosomes showed C9ORF72 puncta on their periphery (Fig. 7B). As expected, starvation led to larger LAMP1-positive lysosomes as a result of membrane fusion (Yu et al., 2010). The localization of C9ORF72 to lysosomes was not obviously altered by serum starvation (Fig. 7B).

**Figure 7:**
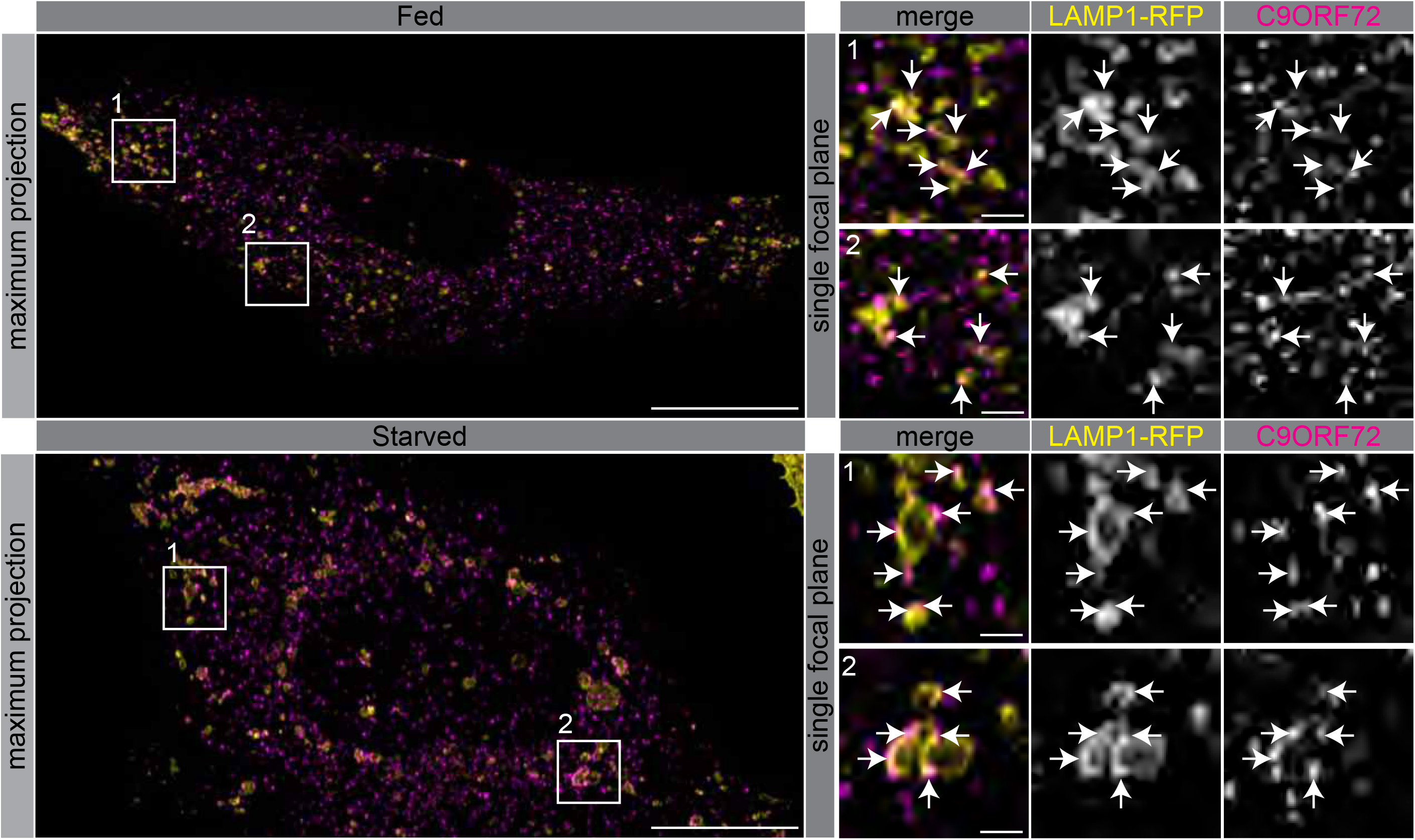
C9ORF72 localizes in part to lysosomes. Immunofluorescence of endogenous C9ORF72 stained with GTX632041 in U2OS cells expressing LAMP1-RFP. The large images on the left show a maximum projection to highlight intracellular structures. Insets 1 and 2 are single focal plane, higher magnification images of the boxed regions. Merge and grayscale images of C9ORF72 and LAMP1-RFP are shown. C9ORF72 localization was observed under basal conditions (top panels) or under starvation for 2 h (bottom panels). Scale bars = 20 μm for large images and 2.0 μm for insets.

Since the highest expression of C9ORF72 we have detected is in macrophages, we sought to investigate localization in these cells. In human MDMs, C9ORF72 decorates large organelles that appear to be phagosomes (Fig. 8A). Macrophages are specialized phagocytic cells that engulf extracellular pathogens and cell debris into phagosomes. Early phagosomes are coated with F-actin (Liebl and Griffiths, 2009) before they mature into late phagosomes that lose their F-actin coat. Late phagosomes than fuse with lysosomes to form phagolysosomes. To confirm that C9ORF72 decorates phagosomes, we preceded with bead-capture experiments. First, RAW264.7 cells were differentiated from monocytes to macrophages by treating them with 1 μg/ml of LPS for 24 hr. Surprisingly, LPS treatment lead to an ~1.8 fold increase in C9ORF72 protein levels (Fig. 8B). We then added 2.3 μm diameter beads and internalization was allowed for 30 min (pulse period) (Fig. 8C). Cells were then washed and were chased (removal of the beads) for the indicated chase period. F-actin was used to label early-phagosomes, whereas LAMP1 was used to label phagolysosomes (Fig. 8C). After a 30 min pulse (30-0), F-actin-coated early phagosomes are devoid of LAMP1, as expected, and devoid of C9ORF72. After a 30 min chase period (30-30), late phagosomes lost the majority of their actin coat, are devoid of LAMP1, but started to accumulate C9ORF72. A 90 min chase period (30-90) allowed enough time for late phagosomes to fuse with lysosomes. Phagolysosomes were devoid of F-actin and accumulated both LAMP1 and C9ORF72 (Fig. 8C). Thus, C9ORF72 starts to accumulate on late-phagosomes once F-actin is being disassembled. C9ORF72 localizes before lysosome fusion on late-phagosomes and remains present on phagolysosomes.

**Figure 8:**
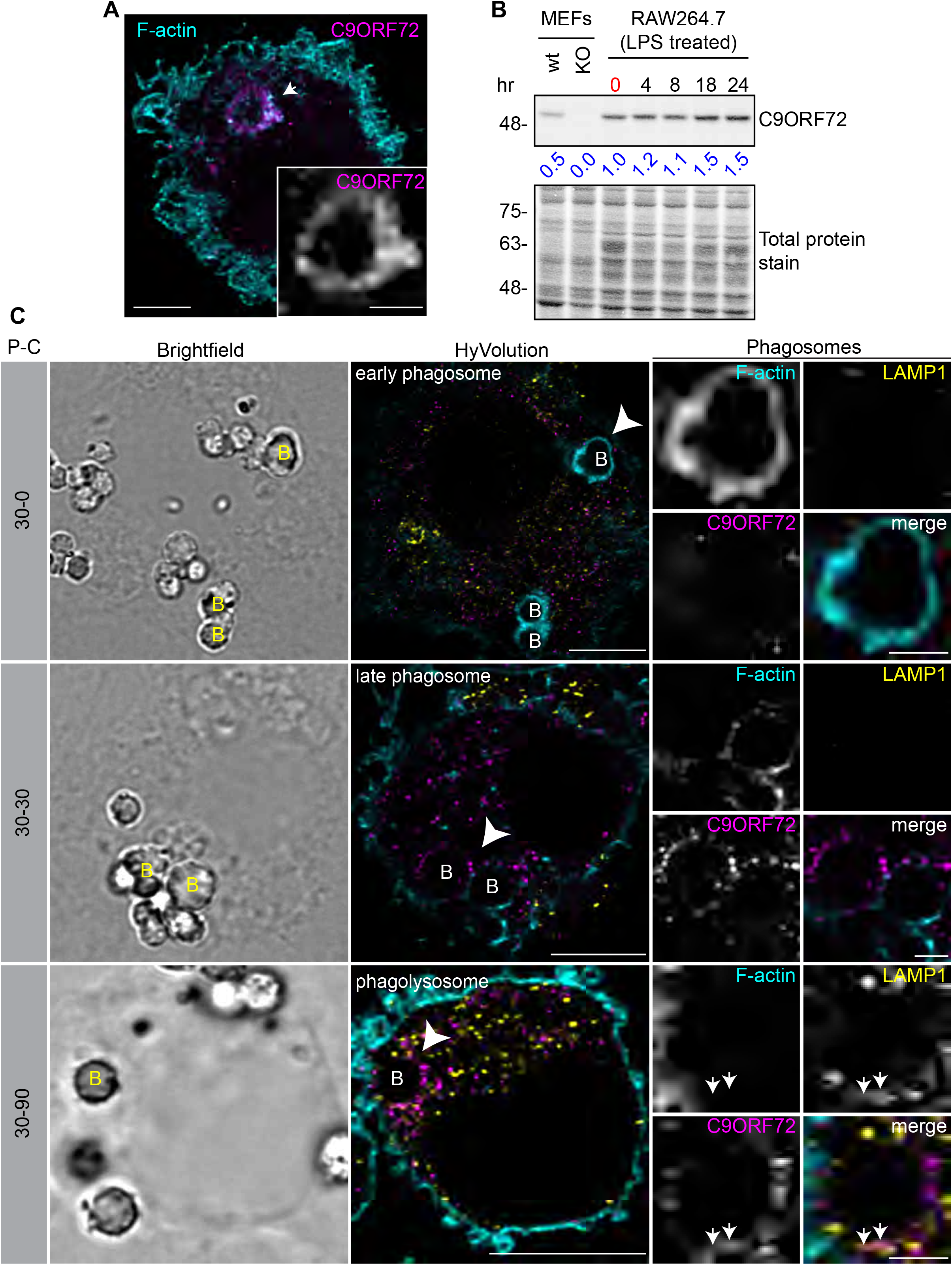
C9ORF72 localizes in part to late phagosomes and phagolysosomes. **(A)** Immunofluorescence of C9ORF72 (GTX632041, magenta) and F-actin staining (cyan) in human MDMs. The inset is a higher magnification of the region indicated by the arrow and shows a grayscale image of C9ORF72. Scale bar = 5 μm (full size) and 2 μm (inset). **(B)** Quantitative immunoblot of RAW264.7 cell lysates treated with 1 μg/ml LPS for the indicated time points. The total protein stained transfer is shown as loading control. C9ORF72 is detected using GTX634482. The C9ORF72 protein signal as a ratio to total protein was determined, normalized to RAW264.7 lysates at time 0 (red), and presented as fold change (blue numbers). C9ORF72 KO MEFs are included as a specificity control. **(C)** Pulse-chase (P-C) experiments in LPS-treated RAW264.7 cells using phagocytosed beads. Beads were incubated for a 30 min pulse followed by a 0 min (30-0), 30 min (30-30) or 90 min (30-90) chase. Brightfield imaging reveals the phagocytosed beads. Staining of C9ORF72 (GTX632041; magenta), LAMP1 (yellow) and F-actin (cyan) were performed and imaged using confocal microscopy together with HyVolution image processing as described previously. Single focal plane images are shown. Insets are higher magnification views of the internalized beads indicated with arrowheads in the large images and show the merged channel together with grayscale images of F-actin, LAMP1 and C9ORF72. White arrows point toward C9ORF72 and LAMP1 colocalization. Scale bar = 8 μm (full size) and 2 μm (inset).

In conclusion, we have developed an antibody validation process that is effective in identifying high-quality antibodies from pre-existing commercial pools and revealing antibodies that should not be used or should be used with extreme caution. With high-quality antibodies new biology can emerge. The technologies employed are simple, easily applicable to any cell biological laboratory, and scalable. Most importantly, implementation of the procedures formalized in this study can serve as a template for granting agencies and journals as a mechanism to ensure enhanced reproducibility.

## Discussion

The scientific community, including academic scientists and commercial antibody producers recognize that there is a crisis in reproducibility caused by the use of non-validated antibodies. The problem could be alleviated with proper characterization of antibodies, but progress to this end has been impeded by technical, economic and sociological challenges. On the technological front, there are numerous types of antibodies (polyclonal, monoclonal, recombinant) used in many different applications - and each combination will require bespoke characterization. It is time-consuming and for some applications there is no quantitative readout. Moreover, these characterization processes are expensive and beyond the budget of any single antibody producer. Finally, there is no consensus among often conflicted scientists and companies, many of whom produce the antibodies, as to what the ideal antibody characterization pipeline should comprise. The advent of CRISPR genome-editing technology provides a powerful new approach to aid in antibody validation, and points to the possibility of creating a more standard antibody characterization approach that depends upon the use of KO controls.

Here we develop a simple, reproducible and relatively inexpensive approach for antibody validation. While this pipeline is for the most part straight forward, there are factors that need to be considered at each step. One important consideration is the choice of a target-expressing cell line to initiate the studies. We used PaxDb, a proteomic database providing protein abundance across multiple tissues/cell types (Wang et al., 2015) (https://pax-db.org/), which indicated that C9ORF72 is expressed in multiple cell lines (HEK-293, HeLa, U2OS) at equal or higher levels than brain, a result confirmed by quantitative immunoblot. In this context, it is important to avoid preconceived notions regarding protein distribution. For example, it is often assumed that neurodegenerative disease genes are predominantly expressed within the neuronal population that is most sensitive to degeneration, yet it is emerging that this is often not the case. Indeed very few are expressed specifically in the nervous system and most are found in multiple cell and tissue types (see the Genotype-Tissue Expression Project; https://gtexportal.org/home/). One example that parallels our findings for C9ORF72 is the major Parkinson disease gene LRRK2 that is broadly expressed in non-neuronal cells, and even within the nervous system, LRRK2 expression levels are low in the most affected area, the substantia nigra (Rui et al., 2018). In fact LRRK2 is most highly expressed in immune cells (Dzamko, 2017). However, PaxDb is limited by the lack of data on some cell types. Our study reveals that the highest levels of C9ORF72 are in macrophages and yet we could not determine this with PaxDb as the database has no proteomic data on this cell type. Therefore, an immunoblot-validated antibody should be used early in the pipeline to screen multiple cell lines.

Another factor to consider are the redundancies seen in commercial antibodies as companies cross-license. For C9ORF72 we identified over 100 commercial antibodies from multiple companies, but when eliminating redundancies, we recognized 14 unique antibodies. While this analysis was tedious, the process could be improved by searches involving Research Resource Identifiers (RRID) that provide unique and permanent identifiers to every antibody (https://scicrunch.org/resources). The number of antibodies available for each target is variable with, for example, 8 antibodies for the newly identified ALS disease gene NEK1 and over 700 for SOD1, an abundant and well-studied ALS gene. If the number of antibodies is unmanageable, it would be advisable to limit validation studies to renewable antibodies such as monoclonals (Andrews et al., 2019). Also, whereas the cost of antibody purchase can appear to be a barrier, many companies have refund policies that can decrease costs if the antibodies to not meet reasonable functional criteria.

Perhaps the most important considerations are the conditions for the analysis by immunoblot, immunoprecipiation, and immunofluorescence. For immunoblot these involve but are not limited to the method of cell lysis, the amount of material analyzed, the type of gel, the transfer method, and the immunoblot conditions. For immunoprecipiation the cell lysis conditions are particularly important as it is vital to generate a fully soluble lysate that extracts the target protein yet maintains native structure. Perhaps the most challenging procedure is immunofluorescence. In our study we were unable to observe a specific signal in HEK-293 cells despite multiple attempts in which we varied fixation and permeabilization conditions. However, when we analyzed U2OS and macrophages, which have higher expression levels, we obtained a specific signal with little effort using 4% PFA followed by Triton X-100 permeabilisation. In a multi-variant screen of fixation conditions for immunofluorescence, it was concluded that 4% PFA followed by Triton X-100 permeabilisation will suffice for the vast majority of targets (Stadler et al., 2010). Therefore, we stress that one needs to perform the immunofluorescence studies in parental versus KO cells after determining an appropriate, highly expressing cell line by immunoblot to ensure a high signal-to-noise ratio (Figure 1). Given that the human proteome contains a vast range of proteins with highly variant chemical compositions, it is difficult to envision set protocols for any of these procedures. It is a challenge to the pipeline to determine empirically the appropriate parameters and this is best left to Investigators with knowledge of selected targets of interest.

Finally, it is important to mention that all antibodies need to be tested under all conditions since the performance and specificity of an antibody depends on the nature of the binding to its protein target, including if the recognition involves linear or conformational epitopes, which must be determined empirically (Mantis et al., 2015). A clear example in this study is GTX634482 that has optimal properties for immunoblot and immunohistochemistry but is not specific in immunofluorescence and is totally negative in immunoprecipitation.

The two GeneTex mouse monoclonal antibodies effective for immunoblot, immunoprecipitation, immunohistochemistry and immunofluorescence have not yet been used in a published paper, whereas the GeneTex rabbit polyclonal antibody (GTX119776), which does not detect the protein by blot and only weakly immunoprecipitates has been used in 5 published papers (Table 1). A troubling finding of our study is that the rabbit polyclonal antibody SC-138763, which clearly does not recognize C9ORF72 in any application, has been used in 15 published manuscripts to ascribe specific properties to the protein in normal and disease states (Table 1). A Google Scholar-based bibliometric approach revealed that those 15 papers have been cited greater than 3000 times and that first layer citations to those citing papers is greater than 66000 citations. While clearly the majority of first and second layer citations do not focus on discoveries related to the use of the antibody in the original 15 papers, this simple analysis reveals the cumulative and sustained impact of inadequate characterization of antibodies.

## Material and Methods

### Antibodies

All C9ORF72 antibodies are listed in Table 1. SMCR8 antibody is from Abcam (ab202283). WDR41 antibody is from Abgent (AP10866B-EV). Peroxidase-conjugated goat anti-mouse and anti-rabbit are from Jackson ImmunoResearch Laboratories. Odyssey IRDye 800CW goat anti-mouse-HRP, the REVERT total protein stain solution and the Odyssey Blocking buffer (TBS) are from LI-COR Biosciences. Alexa Fluor 647-conjugated mouse and rabbit secondary antibodies are from Invitrogen.

### Mouse breeding

Mice were bred and cared for in accordance with the guidelines of the Canadian Council on Animal Care following protocols approved by the University of Toronto Animal Care Committee. C9ORF72 KO mice were a generous gift from Dr. Don Cleveland (UCSD) and Dr. Clothilde Lagier-Tourenne (UMass). In-house breeding and genotyping were performed as previously described (Jiang et al., 2016). Briefly, heterozygous C9ORF72 mice (C57BL/6 background) were crossed to produce homozygous C9ORF72 KO mice, heterozygous C9ORF72 mice, and wild-type littermates. For all experiments, approximately 3-month old C9ORF72 KO and wild-type littermates were used, which was an empirically selected time point. Mouse tissues were prepared for immunohistochemistry or immunoblot as described in separate sections below.

### Cell culture

HEK-293 were cultured in DMEM high-glucose (GE Healthcare cat# SH30081.01) containing 10% bovine calf serum (GE Healthcare cat# SH30072.03), 2 mM L-glutamate (Wisent cat# 609065, 100 IU penicillin and 100μg/ml streptomycin (Wisent cat# 450201). U2OS were cultured in DMEM high-glucose containing 10% tetracyclin-free fetal bovine serum (FBS) (Wisent cat# 081150) 2 mM L-glutamate, 100 IU penicillin and 100 g/ml streptomycin. Tetracyclin-free FBS was used to limit Cas9 expression. U2OS were starved for 2 h in Earle’s balanced salts medium (Sigma cat# E2888).

### CRISPR/Cas9 genome editing

For the KO HEK-293 cells, a C9ORF72 KO gRNA was annealed and ligated into Bbs1-digested pSpCas9(BB)-2A-Puro (PX459) V2.0 vector. PX459 V2.0 was a gift from Feng Zhang (Addgene plasmid # 62988). The gRNA was designed using the “optimized CRISPR design” tool (www.crisp.mit.edu). Oligonucleotides with Bsb1 cleavage overhangs were annealed and cloned into the Cas9/puromycin expressing vector (PX459 from Addgene #48139). The following gRNA sequence was used to create the KO line: CAACAGCTGGAGATGGCGGT. Genome editing plasmids expressing gRNA were transfected in HEK-293 cells using jetPRIME Transfection Reagent (Polyplus) according to the manufacturer’s protocol. The next day, transfected cells were selected using 2 μg/ml puromycin for 24 h. At 96 h post-selection, cells were isolated by clonal dilution. Following the selection expansion of colonies, KOs were confirmed by sequencing of PCR-amplified genomic DNA (data not shown) and immunoblot. Genomic DNA was extracted using QuickExtract DNA extraction solution (Epicentre Biotechnologies).

For the KO U2OS cells, we first generated a line with stable-inducible Cas9. The line was generated using the pAAVS1-PDi-CRISPRn knockin vector and the TALEN pair arms were gifts from Bruce Conklin (Addgene plasmid # 73500) (Mandegar et al., 2016). U2OS cells were resuspended in complete media and 1×10^5^ cells were mixed with pAAVS1-PDi-CRISPRn knockin vector (0.25 ug) and each AAVS1 TALEN pair (0.1 ug) and nucleofected using the Neon® Transfection System (program 17; Life Technologies). Polyclonal population was selected with 2 μg/ml of puromycin (Bioshop cat# PUR555). Seventy-two hours post-treatment, single cells were plated in a 96-well plate. U2OS-PDi-CRISPRn positive clones were detected by PCR and the presence of the PDi-CRISPRn knockin platform into the safe-harbor AAVS1 locus was confirmed by Sanger sequencing.

To generate the C9ORF72 U2OS KO, synthetic single gRNAs (sgRNAs) were designed using the CRISPR Design Tool from Synthego (https://www.synthego.com/products/bioinformatics/crispr-design-tool). The sgRNAs (from Synthego) were transfected in the stable-inducible Cas9 U2OS cell line using JetPRIME transfection reagent. At 1 hr post transfection, Cas9 expression was induced by treating cells with 2 μg/ml of doxycycline (Bioshop cat# DOX44). The next day, media was changed to remove doxycycline and stop the expression of Cas9. C9ORF72 KO cells were isolated by clonal dilution and KOs were confirmed by sequencing of PCR-amplified genomic DNA and immunoblot. Genomic DNA was extracted using QuickExtract DNA extraction solution (Epicentre Biotechnologies). C9ORF72 targeting sgRNA1: GCAACAGCUGGAGAUGGCGG, C9ORF72 targeting sgRNA2: GUCUUGGCAACAGCUGGAGA, AAVS1 locus targeting sgRNA1.2: GGGGCCACUAGGGACAGGAU and AAVS1 locus targeting sgRNA 1.3: GUCCCCUCCACCCCACAGUG

### Immunoblot

Cultured cells were collected in HEPES lysis buffer (20 mM HEPES, 100 mM sodium chloride, 1 mM EDTA, 5% glycerol, 1% Triton X-100, pH 7.4) supplemented with protease inhibitors. Following 30 min on ice, lysates were spun at 238,700xg for 15 min at 4°C and equal protein aliquots of the supernatants were analyzed by SDS/PAGE and immunoblot. Mouse tissues were homogenized in HEPES lysis buffer (without detergent) using a glass/Teflon homogenizer with 10 strokes at 2000 rpm and Triton X-100 was added to 1% final concentration. Following 30 min on ice, mouse tissues lysates were spun at 238,700xg for 15 min at 4°C. Human tissues (including HEK-293 for comparison) were homogenized in RIPA buffer (50mM Tris, 150mM NaCl, 1mM EDTA, 1% sodium deoxycholate, 1% Nonidet P-40, 0,1% sodium dodecyl sulfate, pH 7,4) supplemented with protease inhibitors. Equal protein aliquots of the supernatants were analyzed by SDS/PAGE and immunoblot. Immune cell populations were prepared from spleen, thymus and bone marrow by disrupting the tissue in media using a sterile syringe plunger as a pestle and passing the slurry through a 40 μm cell strainer. Red blood cells from were removed using a red blood cell lysis buffer (155 mM NH_4_Cl, 12 mM NaHCO_3_, 0.1 mM EDTA). Equal protein aliquots of the supernatants were analyzed by SDS/PAGE and immunoblot.

Immunoblots were performed with large 5-16% gradient polyacrylamide gels and nitrocellulose membranes. Proteins on the blots were visualized by Ponceau staining. Blots were blocked with 5% milk, and antibodies were incubated O/N at 4°C with 5% bovine serum albumin in TBS with 0.1% Tween 20 (TBST). The peroxidase conjugated secondary antibody was incubated in a 1:10000 dilution in TBST with 5% milk for 1 h at room temperature followed by washes. For quantitative immunoblot in figure 2, nitrocellulose transfers were incubated with REVERT total protein stain to quantify the amount of protein per lane, than blocked in Odyssey Blocking buffer (TBS) and C9ORF72 antibody GTX634482 was incubated O/N at 4°C in TBS, 5% BSA and 0.2% Tween-20. The secondary antibody (Odyssey IRDye 800CW) was incubated in a 1:20000 dilution in TBST with 5% BSA for 1 h at room temperature followed by washes. Detection of immuno-reactive bands was performed by image scan using a LI-COR Odyssey Imaging System (LI-COR Biosciences) and data analysis was done using LI-COR Image Studio Lite Version 5.2.

### Immunoprecipitation

HEK-293 cells were collected in HEPES lysis buffer supplemented with protease inhibitors. Following 30 min on ice, lysates were spun at 238,700xg for 15 min at 4°C. One ml aliquots at 1 mg/ml of lysate were incubated for ~18 h at 4°C with either 1 μg of a C9ORF72 antibody coupled to protein A or G Sepharose. Beads were subsequently washed four times with 1 ml of HEPES lysis buffer, and processed for SDS-PAGE and immunoblot. For immunoprecipitation with GTX632041 prior to mass spectrometry, HEK-293 cells (parental and C9ORF72 KO) were collected in HEPES lysis buffer supplemented with protease inhibitors. Following 30 min on ice, lysates were spun at 238,700xg for 15 min at 4°C. One ml aliquots at 1 mg/ml were incubated with empty protein G Sepharose beads for 30 min to reduce the levels of proteins bound non-specifically. These pre-cleared supernatants were incubated for 4 h at 4°C with GTX632041 antibody coupled to protein G Sepharose. Following centrifugation, the unbound fractions were collected and the beads were washed 3 times with 1 ml of HEPES lysisi buffer. The beads were then suspended in 1X SDS gel sample buffer. Fractions of the sample were processed for immunoblot. Parallel fractions were run into a single stacking gel band on SDS-PAGE gels to remove detergents and salts. The gel band was reduced with DTT, alkylated with iodoacetic acid and digested with trypsin. Extracted peptides were re-solubilized in 0.1% aqueous formic acid and loaded onto a Thermo Acclaim Pepmap (Thermo, 75 uM ID X 2 cm C18 3 uM beads) precolumn and then onto an Acclaim Pepmap Easyspray (Thermo, 75 uM X 15 cm with 2 uM C18 beads) analytical column separation using a Dionex Ultimate 3000 uHPLC at 220 nl/min with a gradient of 2-35% organic (0.1% formic acid in acetonitrile) over 2 h. Peptides were analyzed using a Thermo Orbitrap Fusion mass spectrometer operating at 120,000 resolution (FWHM in MS1) with HCD sequencing at top speed (15,000 FWHM) of all peptides with a charge of 2+ or greater. The raw data were converted into *.mgf format (Mascot generic format) for searching using the Mascot 2.5.1 search engine (Matrix Science) against Rat protein sequences (Uniprot 2017). The database search results were loaded onto Scaffold Q+ Scaffold_4.4.8 (Proteome Sciences) for statistical treatment and data visualization.

### Immunofluorescence

For antibody validation, U2OS cells (parental and C9ORF72 KO) were transfected with LAMP1-YFP and LAMP1-RFP, respectively. LAMP1-YFP (Addgene plasmid #1816) and LAMP1-RFP (Addgene plasmid #1817) are gifts from Walther Mothes. At 24 h post transfection, both cell lines were plated on glass coverslips as a mosaic and incubated for 24 h. Cells were then fixed in 4% PFA for 10 min, and then washed 3 times. Cells were then blocked and permeabilized in blocking buffer (TBS, 5% BSA and 0.3% Triton X-100, pH 7.4) for 1 h at room temperature. Coverslips were than incubated face down on a 50 μl drop (on paraffin film in a moist chamber) of blocking buffer containing the primary C9ORF72 antibodies diluted at 2 μg/ml and incubated overnight at 4°C. Cells were washed 3 x 10 min and incubated with corresponding Alexa Fluor 647-conjugated secondary antibodies diluted 1:1000 in blocking buffer for 2 h at room temperature. Cells were washed 3 x 10 min with blocking buffer and once with TBS. Coverslips were mounted on a microscopic slide using fluorescence mounting media (DAKO, Cat# S3023). HEK-293 (parental and C9ORF72 KO) were plated separately on coverslips and stained as mentioned above. HEK-293 cell lines were fixed in either 4% PFA for 10 min or in methanol (chilled at −20°C) for 10 min. Imaging was performed using a Leica SP8 laser scanning confocal microscope equipped with a 40x oil objective (NA = 1.30) and HyD detectors. Acquisition was performed using Leica Application Suite X software (version 3.1.5.16308) and analysis was done using Image J. All cell images represent a single focal plane. They were prepared for publication using Adobe Photoshop to adjust contrast, apply 1 pixel Gaussian blur and then assembled with Adobe Illustrator.

Imaging of C9ORF72 in U2OS cells at higher resolution was done by transfecting LAMP1-RFP and C9ORF72 staining was performed as above. Imaging was performed using a 63x oil objective (NA = 1.40) and we used HyVolution (Leica) to acquire images using the best setting for the resolution of the deconvolved image.

### Immunohistochemistry with 3,3’-diaminobenzidine (DAB) staining

Formalin-fixed, paraffin-embedded brain tissue from C9-KO (n=3) and C9-wild-type (n=3) mice were sectioned at 6 μm in the sagittal plane and then mounted onto positively charged slides. Sections were deparaffinised at 60°C for 20 min on a heat block and then incubated in xylene (3 x 5 min). Subsequent rehydration was performed through graded ethanol washes and finally in water. For epitope unmasking, heat-induced epitope retrieval was carried out using TE9 buffer (10 mM Trizma base, 1 mM EDTA, 0.1% Tween 20, pH 9 at 110°C for 15 min. Endogenous peroxidases were quenched with 3% H_2_O_2_ in TBS for 10 min at room temperature. Slides were then blocked in 2.5% normal horse serum and 0.3% Triton X-100 in TBS for 1 h at room temperature. GTX63448 and GTX632041 were diluted in DAKO antibody diluent (Agilent, Cat# S0809) at 1:5000 and then slides were incubated overnight at 4°C. Washes were then performed 3 x 10 min in TBS-0.1% Tween 20 (TBST) prior to secondary antibody incubation with ImmPRESS HRP horse anti-mouse IgG (Vector Labs, Cat# MP-7402), for 1 h at room temperature. Slides were then washed 3 x 20 min in TBST. DAB staining was developed under a light microscope for between 2-10 min as per the manufacturer’s instructions with the ImmPACT DAB peroxidase substrate kit (Vector Labs, Cat# SK-4105). Slides were then counterstained with Hematoxylin Solution, Gill No.1 (Sigma-Aldrich, Cat# GHS132) for 5 min at room temperature. Slides were then dehydrated with sequentially increasing graded ethanol and finally in xylene prior to coverslipping with Cytoseal 60 (Fisher Scientific, Cat# 8310-16). Micrographs were captured with a digital camera mounted on a Leica DM6000 B upright microscope with either 10X or 40X objectives using Volocity Imaging software (version 6.3.0 PerkinElmer).

### Rapid immune-isolation of lysosomes (LysoIP)

The isolation of lysosome was performed following the LysoIP protocol (Abu-Remaileh et al., 2017) with slight modifications. Both Tmem192-3xHA (Addgene #102930) and Tmem192-2xFlag (Addgene #102929) were gifts from David Sabatini. Cells were transfected with either Tmem192-3xHA (HA-Lyso cells) or with TMEM192-2xFlag (Control-Lyso cells). Around 35 million cells were used for each LysoIP. Cells were quickly rinsed twice with cold PBS and scraped in 2 ml of KPBS (136 mM KCl, 10 mM KH_2_PO_4_, pH 7.25 and protease inhibitor) and centrifuged at 1000 x g for 2 min at 4°C. Pelleted cells were resuspended in 900μl of KPBS and gently homogenized with 20 strokes in a 2 ml homogenizer. The homogenate was centrifuged at 1000 x g for 2 min at 4°C. The supernatant was collected and 10 μl (equivalent to 1,1%) was reserved and run on immunoblot as the starting material. The remaining supernatant was incubated with 150 μl of KPBS prewashed anti-HA magnetic beads (Thermo Fisher Scientific cat# 88837) on a gentle rotator shaker for 3 min. Beads were gently washed 3 times with 1 ml of KPBS using a DynaMag-2 magnet (Thermo Fisher Scientific cat# 12321D). Beads were resuspended in 1 ml of KPBS and transfer to a new tube. KPBS was removed and beads were incubated in resuspension buffer (20mM Hepes, 1mM EDTA, 150 mM NaCl, 1% Triton X-100, pH 7,4 + phosphatase inhibitor). Supernatant was collected and run on an immunoblot.

### Human tissues isolation

#### Whole brain

Whole brain tissue was taken from surgical resections of brain tissue from pharmacologically intractable, non-malignant cases of temporal lobe epilepsy.

#### Primary Human Microglia

Human microglia were isolated from adult brain tissue using previously described protocols (Durafourt et al., 2013). Briefly, adult microglia were derived from surgical resections of brain tissue from pharmacologically intractable non-malignant cases of temporal lobe epilepsy. Brain tissue was mechanically dissociated and enzymatically digested using trypsin and DNAse prior to mechanical separation through a nylon mesh filter. Tissues then underwent ultracentrifugation to remove myelin. Dissociated cells were centrifuged, counted, and plated at 2-3 x 10^6^ cells/ml in MEM supplemented with 5% FBS and 0.1% penicillin/ streptomycin, and 0.1% glutamine. Microglia were grown for 3 days, collected, plated at 1 x 10^5^ cells/ml, and maintained in culture for 6 days.

#### Human Fetal Astrocytes

Human CNS fetal tissue (cerebral hemispheres) was obtained at 17–23 week of gestation from the Human Fetal Tissue Repository (Albert Einstein College of Medicine, Bronx, NY), following CIHR-approved guidelines. As previously described (Jack et al., 2005), astrocyte cultures were obtained by dissociation of the fetal CNS with trypsin and DNase followed by mechanical dissociation. After washing, the cell suspension was plated at a concentration of 3–5 x 10^6^ cells/ml on poly-L-lysine coated flasks in DMEM supplemented with 10% FCS, antibiotics, glutamine. To obtain pure astrocytes, the mixed CNS cell culture (containing astrocytes, neurons, and microglia) was passaged upon confluency, starting at 2 weeks, post-isolation, using trypsin-EDTA (Invitrogen Life Technologies). Human fetal astrocytes were used between passages 2 and 4 and cultures were determined to be >90% pure, as determined by immunostaining for glial fibrillary acidic protein (GFAP). Astrocytes were treated with 100ng/ml of IL-1β overnight.

#### Monocyte-derived Macrophages

Peripheral blood mononuclear cells (PBMCs) were isolated from whole blood using Ficoll– Paque density gradient centrifugation (GE Healthcare). CD14+ cell isolation was done using immunomagnetic bead selection according to manufacturer’s instructions to achieve 95–99% purity (Miltenyi Biotec). CD14+ monocytes were cultured in RPMI cell medium supplemented with 10% FCS, 0.1% penicillin/streptomycin and 0.1% glutamine. MDMs were differentiated in vitro with 25 ng/ml M-CSF for 6 d, during which time cells they were either activated with the pro-inflammatory cytokines IFN-γ and LPS or the anti-inflammatory cytokine TGF-β, as previously described (Healy et al., 2016). Briefly TGF-β-treated cells received recombinant human TGF-β (20 ng/ml) on days 1 and 4. Pro-inflammatory activated cells received IFN-γ (20 ng/ml) for 1 h followed by a 48 h treatment with LPS (serotype 0127:B8, 100 ng/ml).

All studies with human tissue following Canadian Institutes of Health Research approved guidelines. Secondary use of de-identified tissues was carried out in accordance with the guidelines set by the McGill University Institutional Review Board (McGill University Health Centre Ethics Board and approved under protocol ANTJ1988/89). All experiments were conducted in accordance with the Helsinki Declaration with sample procured with informed consent.

## Acknowledgements

We thank Dr. Don Cleveland (UCSD) and Dr. Clothilde Lagier-Tourenne (UMass) for C9ORF72 KO mice. Proteomics analysis was performed at the Research Institute of the McGill University Health Centre Clinical Proteomics Platform. This work was supported by a grant from the Motor Neurone Disease Association (UK), The ALS Association (USA) and ALS Canada and by an Arthur J Hudson Team Grant from ALS Canada/Brain Canada. CL is supported by the Ronald Peter Griggs and Tim E Noël Postdoctoral Fellowship from ALS Canada. RK is supported by a studentship from the Canada First Research Excellence Fund, awarded to McGill University for Healthy Brains for Healthy Lives. JR holds the James Hunter and Family Chair in ALS Research. PSM is a James McGill Professor and a Fellow of the Royal Society of Canada.

---

**Supplementary Figure 1:**
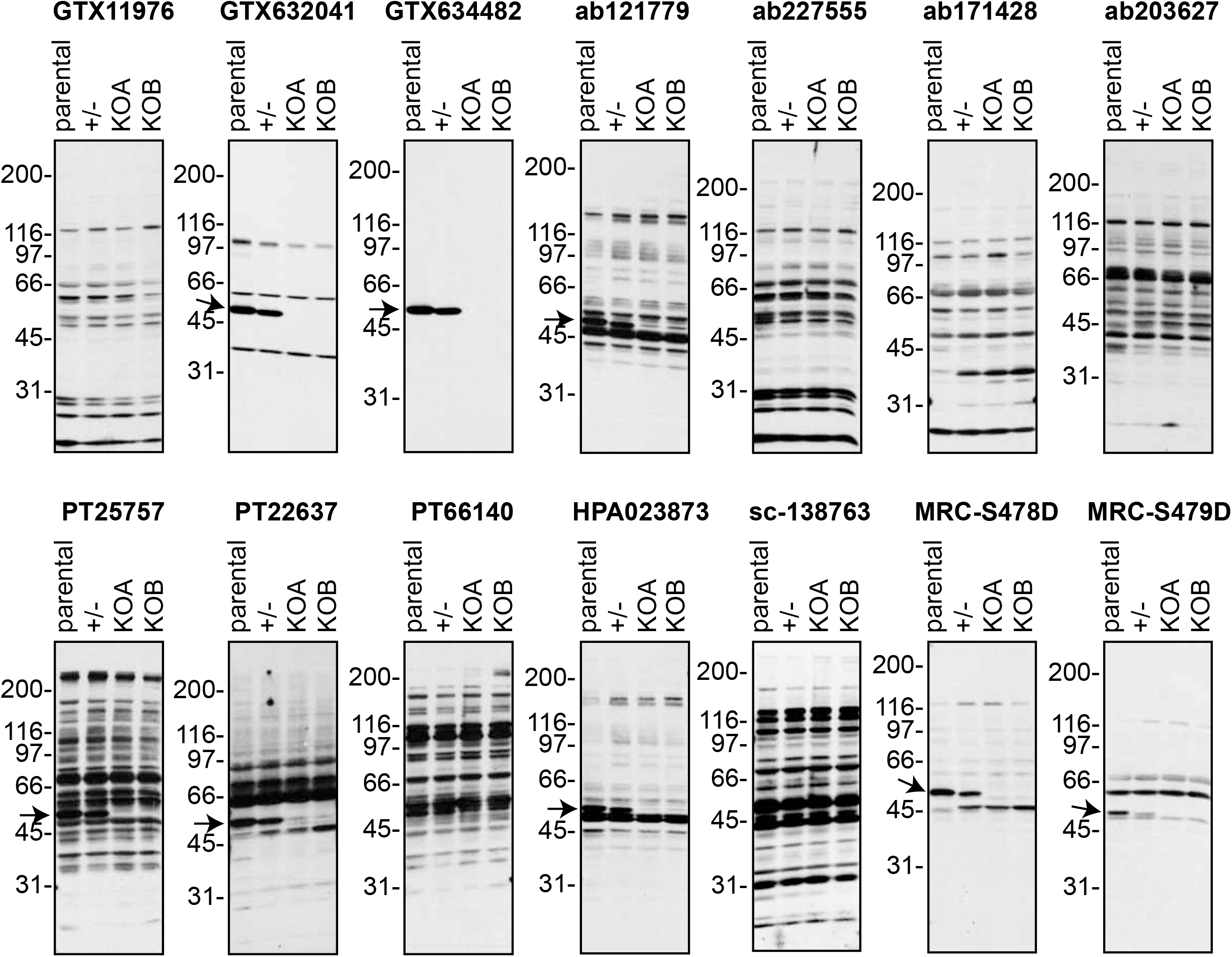
Validation of C9ORF72 antibodies by immunoblot. Cell lysates from HEK-293 parental, heterozygous (+/-) or two individual C9ORF72 KO clones (KOA, KOB) were prepared and processed for immunoblot with the indicated C9ORF72 antibodies. The arrows point to positive C9ORF72 signals.

**Supplementary Figure 2:**
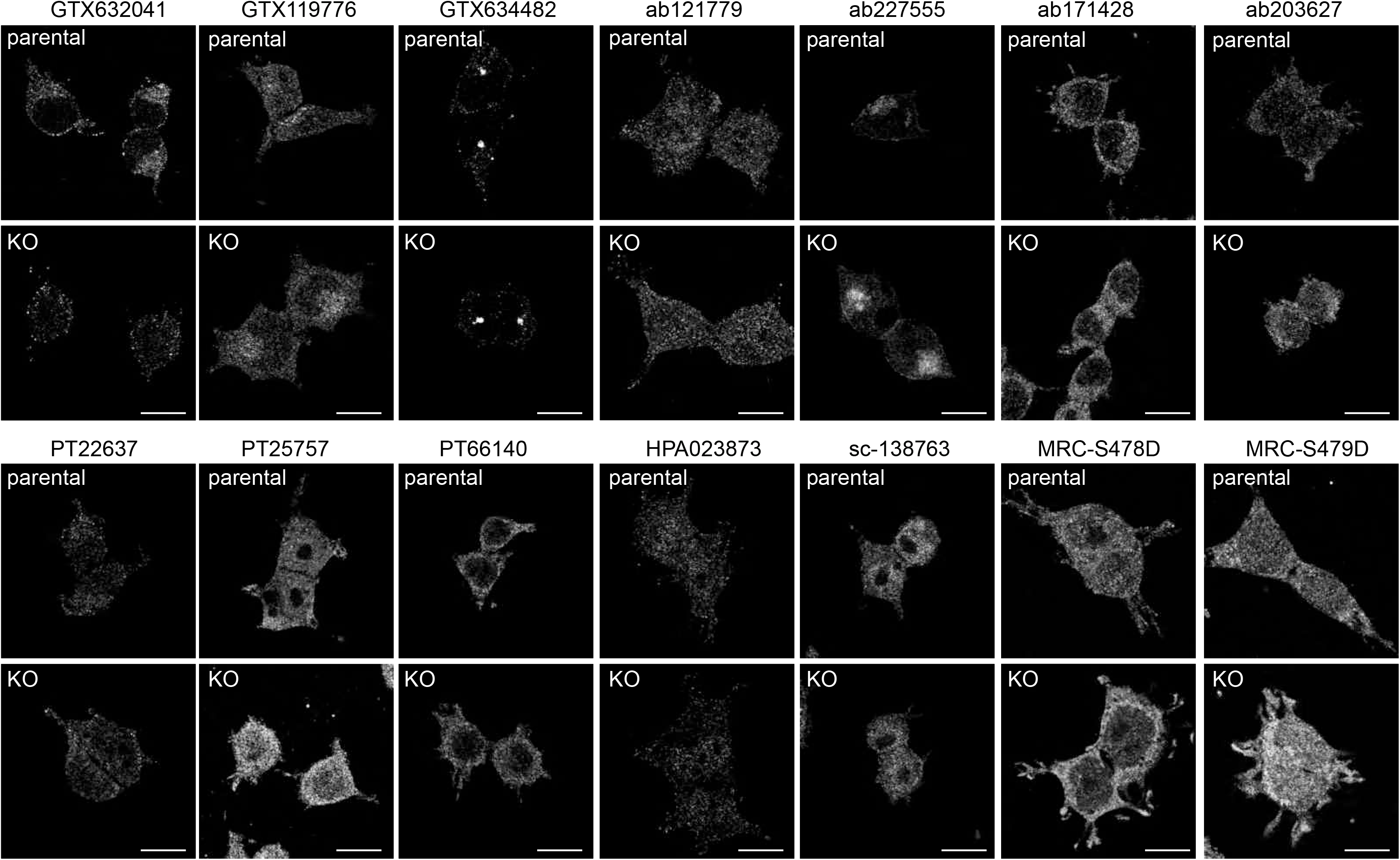
Testing of C9ORF72 antibodies by immunofluorescence analysis in HEK-293 cells fixed with PFA. HEK-293 parental and KO cells were fixed with PFA, permeabilized and incubated with the indicated C9ORF72 antibodies. Representative images are shown. Bars = 16μm.

**Supplementary Figure 3:**
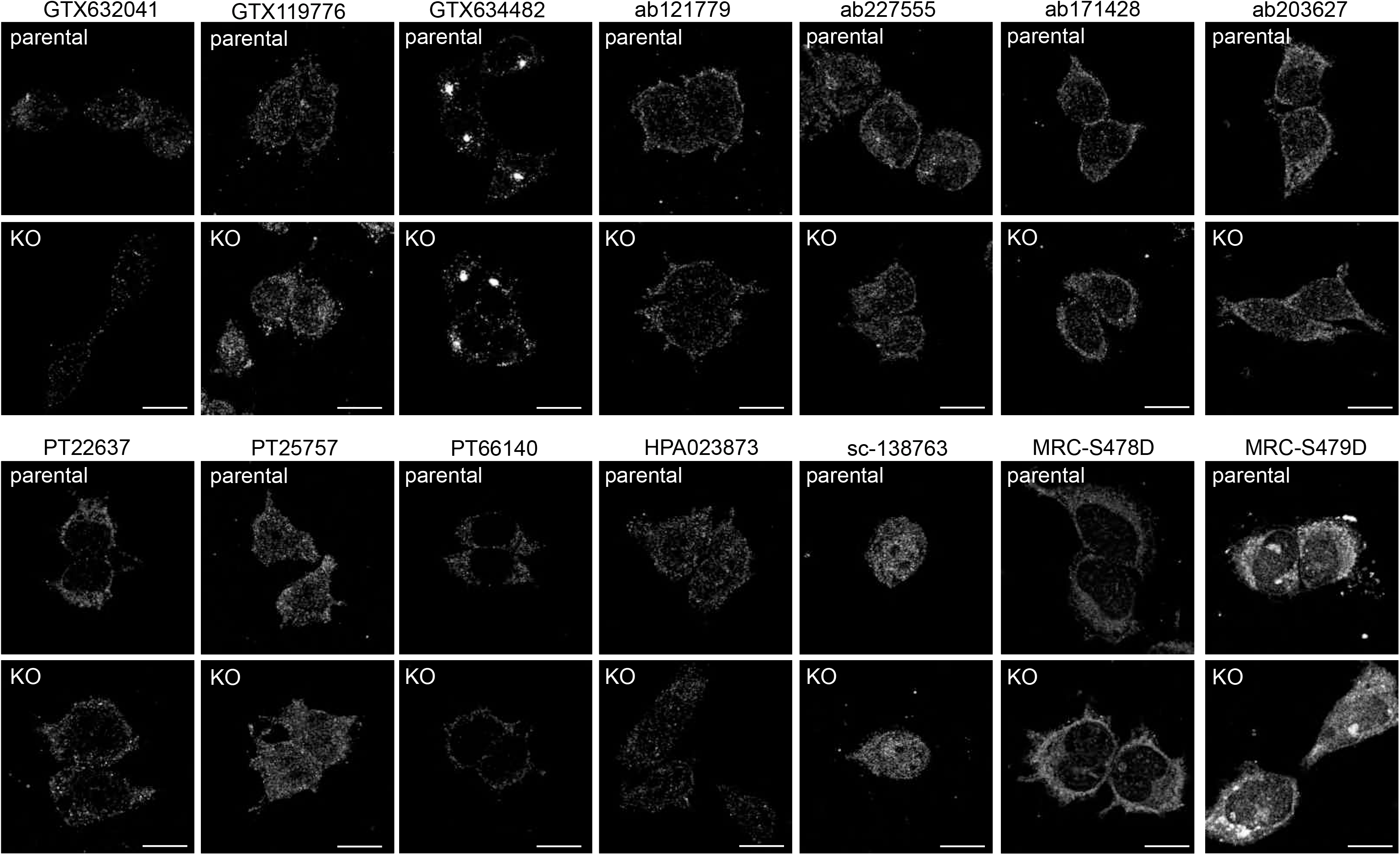
Testing of C9ORF72 antibodies by immunofluorescence analysis in HEK-293 cells fixed with −20°C methanol. HEK-293 parental and KO cells were fixed with −20°C methanol and incubated with the indicated C9ORF72 antibodies. Representative images are shown. Bars = 16μm.

**Supplementary Figure 4:**
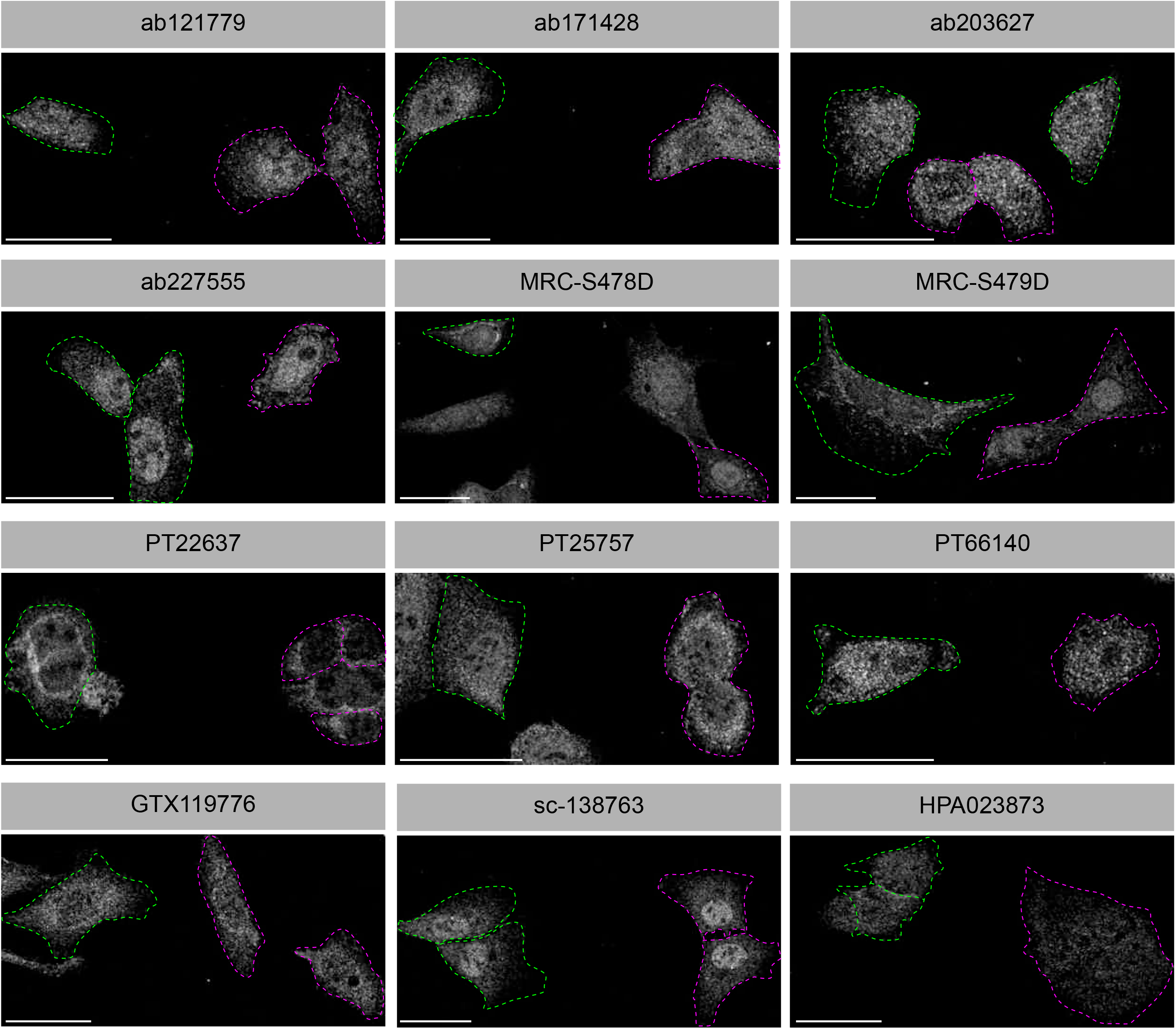
Testing of 12 C9ORF72 antibodies by immunofluorescence analysis. Parental and KO cells were transfected with LAMP1-YFP or LAMP1-RFP, respectively. Parental and KO cells were combined and stained with the indicated C9ORF72 antibodies. Greyscale images of the far-red channel are shown. Parental and KO cells are outlined with green and red dashed line, respectively. Representative images are shown. Bars = 40μm.

**Supplementary Figure 5:**
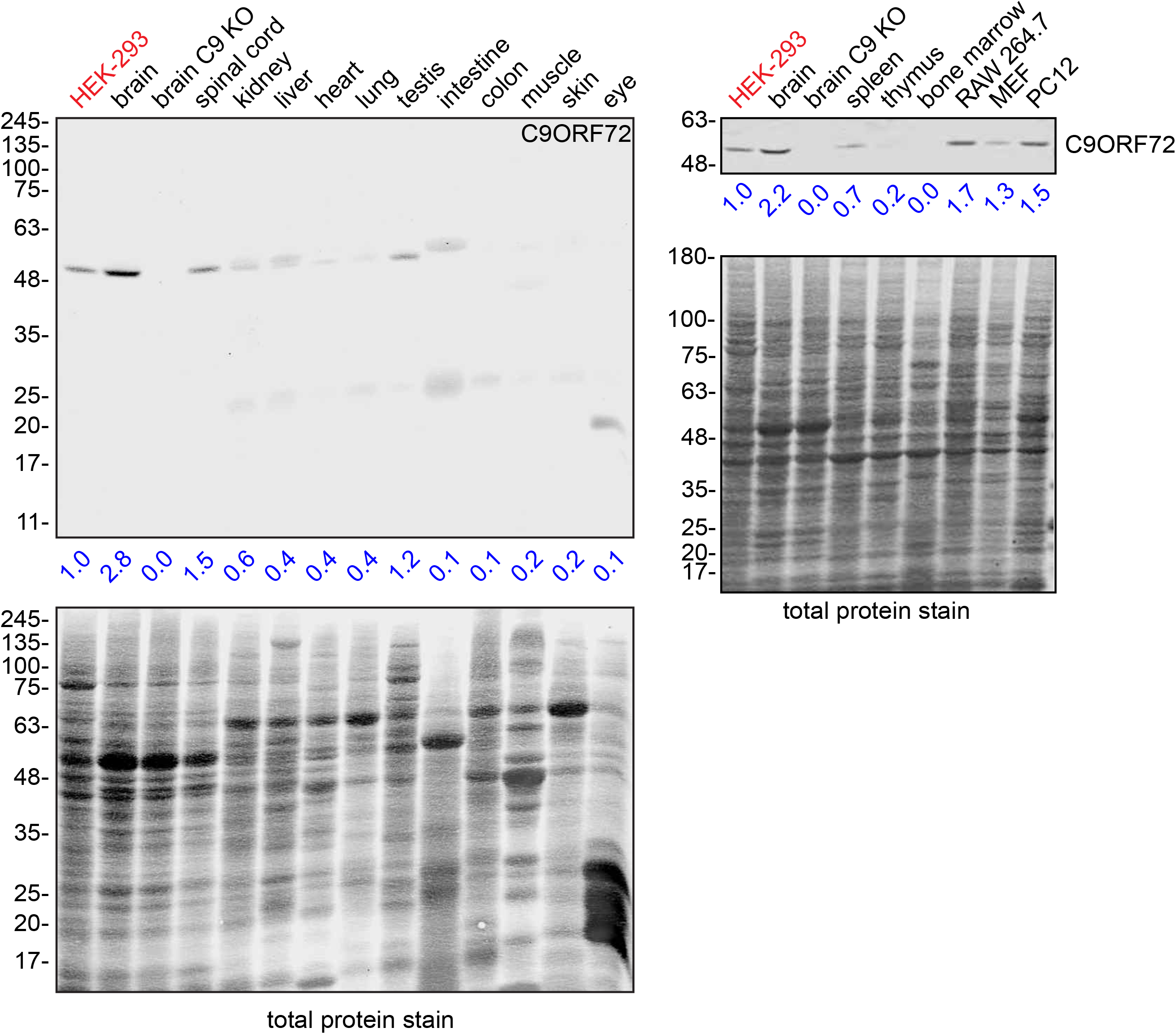
Comparison of C9ORF72 protein level in cell lines and mouse tissues. Lysates were prepared from mouse tissues and selective murine cell lines and processed for quantitative immunoblot using antibody GTX634482 with the LI-COR Odyssey Imaging System as described previously (RAW264.7 = mouse macrophage; MEF = mouse embryonic fibroblast; PC12 = rat pheochromocytoma). The total protein stained transfers are shown as loading controls. The C9ORF72 protein signal as a ratio to total protein was determined, normalized to parental HEK-293 cells (red), and presented as fold change (blue numbers). KO mouse brain (brain C9 KO) was included for specificity.

**Supplementary Figure 6:**
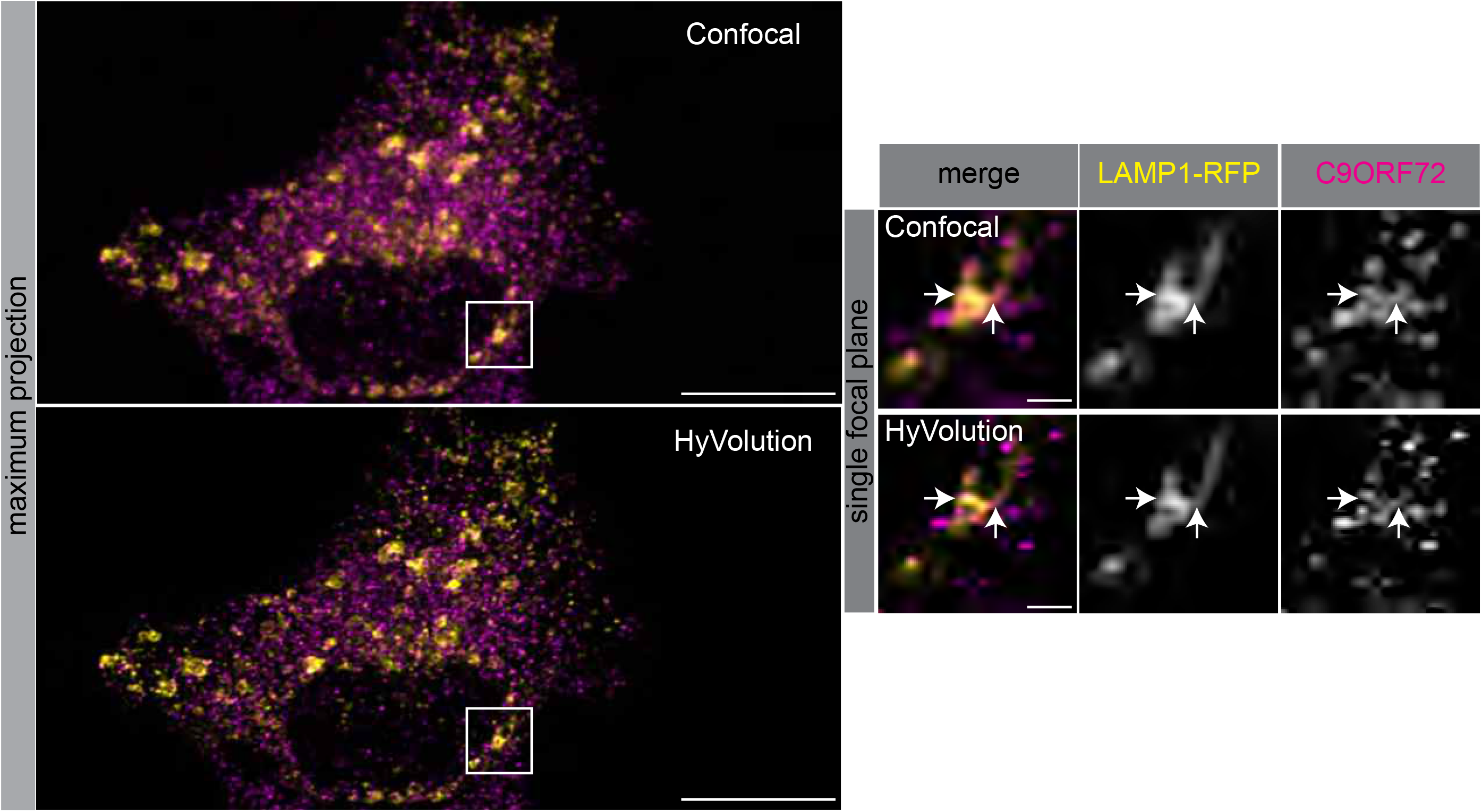
HyVolution enhances confocal image resolution. Immunofluorescence of endogenous C9ORF72 stained with GTX632041 in U2OS cells expressing LAMP1-RFP. Full-size cell image shows a maximum projection. Insets are higher magnification of the boxed regions (single focal plane). Merge and grayscale images of C9ORF72 and LAMP1 are shown. Top panel shows regular confocal imaging. Bottom panel shows the result of HyVolution applied to the above confocal image. Full-size cells bars = 20μm, insets bars = 2.0μm.

